# The crop pathogen *Blumeria hordei* exhibits genome-wide pervasive selective and neutral sweepstakes reproduction signatures

**DOI:** 10.64898/2026.05.05.723056

**Authors:** Miles Anderson, Luzie U. Wingen, Bernarda Biggeman Troche, Xinyi Liu, Marion M. Müller, Ralph Hückelhoven, Aurélien Tellier

## Abstract

The fungal crop pathogen *Blumeria hordei*, causal agent of powdery mildew on barley, presents life-history and epidemiological characteristics, as well as and selective pressures due to modern agriculture leading to expected sweepstakes reproduction, that is highly skewed offspring distributions. Using genome-wide polymorphism data and population genomics inferences, we aim to 1) infer the past demographic history and the strength of sweepstakes reproduction in *B. hordei*, and 2) quantify the contributions of these selective and neutral processes in the genome. An new inference method based on Neural Posterior Estimation and diversity and linkage disequilibrium statistics was developed and tested on simulated and *B. hordei* genomic data. We confirm that B. hordei exhibits a moderate sweepstakes reproduction (*α*-parameter of 1.6). We highlight that the Site Frequency Spectrum (SFS) appears sensitive to the joint occurrence of sweepstakes and recent demographic changes, which may caution on the reliability of the SFS to infer sweepstakes reproduction. We then scan the genome for selective sweeps, adjusting the significance thresholds of the methods for demographic history and sweepstakes reproduction, thereby yielding a counterintuitive result. When conditioning the significance threshold for sweep detection on simulations under sweepstakes and demography, a very large number of putatively selected regions is found (11.6% of the genome). We suggest that sweepstakes reproduction in *B. hordei* is due to 1) neutrality (clonal/sexual phases and Boom-and-Bust cycles) generating a genome-wide level of background noise in the coalescent genealogies, and 2) selective sweepstakes due to pervasive positive selection. Our findings have important implications for both population genomic methodology and our understanding of pathogen evolution.

## 1 Introduction

Widely used traditional population genetic models assume that reproducing individuals exhibit a small variance in offspring production, and as a consequence, that the population genealogies follow the standard (Kingman) coalescent process. The latter is characterized by binary mergers whereby lineages coalesce pairwise moving backward in time (Kingman 1982; Wakeley 2009). This framework has proven invaluable for understanding evolutionary dynamics and past demographic history in many organisms, including the human species. However, the key assumption of small variance in offspring production between parents is violated in many species due to two possible sweepstakes reproduction mechanisms (Schweinsberg 2003; Eldon 2020): 1) selective events, and/or 2) neutral biological and ecological processes. The detection of positive selection events, the so-called selective sweeps (Stephan 2019), in most species, including humans, builds on the shape of the (sample) genealogy to be distorted at the loci under selection due to the high variance in offspring production by selected individuals (Schweinsberg 2003; Durrett and Schweinsberg 2004). This can also be described as multiple merger coalescent (MMC) signatures, allowing for multiple lineages to merge simultaneously during a coalescence event. This mechanism is called selective sweepstakes reproduction. A second, non-mutually exclusive mechanism generating large variance in offspring production arises from neutral processes imposed by biological and ecological life-cycles and life-history traits. Large variance in offspring production can thus be a life-history trait and reproduction strategy of the organism, such as found in fish (Waples 2016; Waples et al. 2018; Árnason et al. 2023), or due to severe recurrent bottlenecks and/or alternating modes of reproduction (Waples et al. 2013; Tellier and Lemaire 2014; Matuszewski et al. 2018; Eldon 2020; Menardo, Gagneux, and Freund 2021). Sweepstakes reproduction can fundamentally alter evolutionary dynamics and polymorphism signals, challenging standard population genetic inference (Sagitov 1999; Schweinsberg 2003; Eldon 2020; Korfmann et al. 2024a; Korfmann et al. 2024b; Goldberg 2026). Under MMC processes, coalescent timescales differ fundamentally from standard Kingman expectations, potentially leading to systematic biases in demographic inference and false signals of selection when traditional methods are applied (Eldon et al. 2015; Montano 2016; Matuszewski et al. 2018; Menardo, Gagneux, and Freund 2021; Freund et al. 2023; Goldberg 2026). MMC footprints have been identified in economically important fish species (Waples et al. 2013; Árnason et al. 2023) and in several invertebrate, protozoan, fungal, bacterial, and viral species, and seem to be especially important characteristics of parasites/pathogens of animals, plants and humans (Montano 2016; Matuszewski et al. 2018; Menardo, Gagneux, and Freund 2021; Freund et al. 2023; Jigisha et al. 2025; Goldberg 2026). Therefore, it may be relevant to dissect and infer the causes and consequences of sweepstakes reproduction to better understand and predict the evolutionary potential of these species. However, the respective importance and contributions of neutral and/or sweepstakes reproduction in generating the MMC signatures have not yet been assessed (Matuszewski et al. 2018; Menardo, Gagneux, and Freund 2021; Árnason et al. 2023; Freund et al. 2023; Jigisha et al. 2025).

A critical limitation of current approaches for detecting and inferring the strength of MMC is their susceptibility to being confounded by demographic history, as well as to confound demographic history inference (Koskela 2018; Koskela and Wilke Berenguer 2019; Freund and Siri-Jégousse 2021; Korfmann et al. 2024b). Indeed, complex population dynamics can generate signatures that mimic or obscure genuine multiple-merger processes (e.g., Matuszewski et al. 2018; Korfmann et al. 2024b), especially when using the site-frequency spectrum (SFS) as a statistic of choice to perform MMC footprint inference (Koskela 2018; Koskela and Wilke Berenguer 2019; Freund et al. 2023; Árnason et al. 2023). The lack of power and limitation of the SFS-based approaches is reflected in several inconclusive analyses regarding the causal evolutionary (neutral versus selective) mechanisms generating MMC footprints (Árnason et al. 2023; Freund et al. 2023; Jigisha et al. 2025). This stems from these methods forcing a choice between demographic or MMC inference. Furthermore, existing SFS-based inference methods do not explicitly model recombination and assume independent sites (e.g., Koskela 2018; Koskela and Berenguer 2019), thereby failing to exploit the power of genealogical patterns along the genome (but see the ancestral recombination graph in Birkner, Blath, and Eldon 2013 and the methods in Korfmann et al. 2024b). To bridge this gap, we aim to build an inference method that improves on Korfmann et al. 2024b to account for the influence of possibly complex demographic effects under MMC signatures. A key signature of MMC with recombination is the possibility of long-range Linkage Disequilibrium (LD) appearing between sites (or coalescent trees) at different places in the genome because they originate from the same MMC event. Such signatures are not expected under the Kingman coalescent model (Birkner, Blath, and Eldon 2013; Korfmann et al. 2024b). In this study, we have two aims. First, we infer the demographic history of species exhibiting MMC footprints, and concomitantly, the parameters of sweepstakes reproduction. Second, we aim to quantify the contributions of selection and neutrality to produce sweepstakes reproduction signatures.

Our motivation stems from the study of the barley (*Hordeum vulgare* L.) system and its pathogen *Blumeria hordei* (*B. hordei*), the causal agent of powdery mildew in barley. Barley represents one of the world’s most important cereal crops, serving critical roles in food security, livestock feed, and malting industries globally (Arendt and Zannini 2013; Kant, Amrapali, and Babu 2016; Stanca et al. 2016). However, barley production faces persistent challenges from numerous fungal pathogens, among which powdery mildew caused by *B. hordei*, although some durable resistance exists (Dreiseitl 2024). *B. hordei* is an obligate biotrophic fungus that exclusively parasitizes barley, forming characteristic white, powdery colonies on leaf surfaces, stems, and inflorescences (Spanu and Panstruga 2012). The pathogen exhibits remarkable evolutionary adaptability, consistently overcoming host resistance genes and fungicidal control (Brown and Jessop 1995; Frantzeskakis et al. 2018; Ellwood, Lopez-Ruiz, and Tan 2024). This evolutionary arms race between the barley host (under human influence) and the pathogen has profound implications for sustainable disease management strategies and necessitates a deeper understanding of the population-genetic processes underlying pathogen evolution.

The evolutionary characteristics of *B. hordei* present significant but underappreciated challenges for traditional population genomic approaches. First, both *Blumeria* species exhibit a high proportion (more than 70%) of Transposable Elements (TEs) in their genome (Spanu et al. 2010; Bindschedler, Panstruga, and Spanu 2016; Frantzeskakis et al. 2018; Liu et al. 2026) which likely have a direct and indirect selective influence on neighboring genes (Frantzeskakis et al. 2018; Liu et al. 2026), and on potential past demographic inference (see the effect of TEs on Sequentially Markovian Coalescent inference, Sellinger, Abu-Awad, and Tellier 2021). Second, the sexual reproduction system involves the formation of chasmothecia containing ascospores, which serve as the primary inoculum for spring infections. The pathogen also reproduces asexually through conidia production during the growing season, creating complex population dynamics with both sexual and asexual phases (Agrios 2024). Critically, these reproductive characteristics may generate highly skewed offspring distributions, whereby a small number of individuals may disproportionately contribute to the next generation (so-called MMC event). Third, *Blumeria* species exhibit epidemic growth patterns and recurrent bottlenecks associated with seasonal cycles (Boom-and-Bust cycles, Agrios 2024) and patchy host availability (see the *oases in the desert* metaphor, Brown et al. 2002). Fourth, in modern agriculture, crop pathogens are under selective pressure to adapt and overcome major resistance genes introduced in new varieties, and to resist applied fungicides (Frantzeskakis, Kusch, and Panstruga 2018). Altogether, these biological and life-history traits create conditions that may favor MMC events (Tellier and Lemaire 2014). Recent evidence for MMC has emerged in *Blumeria graminis* forma specialis *tritici* (*Bgt*), the causal agent of wheat powdery mildew (Jigisha et al. 2025). Given the close taxonomic relationship between *Bgt* and *B. hordei*, as well as their similar life history traits, environmental conditions, and geographical distributions, there is compelling reason to hypothesize that MMC processes may also shape *B. hordei* evolution. However, the extent to which multiple-merger processes influence *B. hordei* population dynamics, and their implications for detecting genuine selective signatures, remain unexplored. For crop pathogens in general, and *B. hordei* in particular, neutral-sweepstakes reproduction is predicted to occur due to regular boom-and-bust cycles and/or alternating clonal/sexual phases, while selective sweepstakes reproduction may be driven by adaptation to crop resistance and fungicide resistance (Tellier and Lemaire 2014).

To overcome the highlighted methodological challenges and enable robust MMC inference in organisms with complex evolutionary histories, we develop a novel approach based on Neural Posterior Estimation (NPE) for inferring coalescent parameters in the presence of arbitrarily complex demographic scenarios. Unlike existing methods, our NPE framework explicitly accounts for recombination and can distinguish MMC signatures from those generated solely by demographic processes. This methodological advance builds on accounting for long-range LD and thus represents a significant step forward in coalescent inference, enabling more accurate characterization of genealogical processes in populations with realistic demographic complexities. Using a dataset of 43 whole genome sequences of *B. hordei* collected within Germany, we test for evidence of multiple-merger genealogies in an endemic population. We characterize the extent to which demographic complexity can confound MMC inference methods based upon the site frequency spectrum (SFS), demonstrate the ability of our NPE framework to infer the MMC parameter in the presence of complex demographic histories, and develop robust statistical assessment of the proportion of adaptive loci. We suggest that 1) the population size of *B. hordei* tracks the usage of Barley in Europe since ca. 1,000 years, and 2) genome-wide signatures of MMC are likely due to selective processes.

## 2 Material and Methods

We investigated the population genomic signatures of *B. hordei* (hereafter *B. hordei*) through the lens of MMC theory, specifically focusing on the *β*-coalescent framework. The *β*-coalescent represents a tractable sub-class of the *λ*-coalescent where the frequency and strength of multiple-merger events are characterized by the parameter *α* ∈ (1, 2] (Schweinsberg 2003; Birkner, Blath, and Eldon 2013. In this model, *α* = 1 represents an extreme MMC process with frequent large merger events, while *α* = 2 approaches the standard Kingman coalescent of binary mergers (Schweinsberg 2003; Eldon et al. 2015; Birkner, Blath, and Eldon 2013; Koskela 2018; Eldon 2020; Freund et al. 2023; Korfmann et al. 2024b).

### 2.1 Sampling and Sequencing of the German *B. hordei* population

We collected 53 *B. hordei* isolates from barley plants in the German federal states of Bavaria and Brandenburg in the years 2023 and 2024 (Figure S1, Table S1). Single-spore purification was conducted for all isolates as described in Liu et al. 2026 for the reference isolate TUM1. Whole genome DNA was extracted from conidiospores as described in Liu et al. 2026 for the reference isolate TUM1. DNA of 53 isolates was used for short-read paired-end sequencing with Illumina NovaSeq™ X Plus (2 x 150 bp) technology (GENEWIZ Germany GmbH: 13 isolates (Bay43, Bay50, Bay52, EL01 01, EL01 02, EL02 02, EL03 01, FS02, FS03, FS04, FS05, FS07, and Muc01) and by Novogen Germany GmbH: all other 30 isolates). Additionally, long-read sequence reads were obtained for 13 isolates using Oxford Nanopore Technology (LMU Gene Centre, Munich) as described in Liu et al. 2026 (Table S1). For two isolates the long-read sequencing reactions delivered only low output.

### 2.2 Variant Calling

The raw Illumina reads were filtered and adapters were removed using BBDuk from the BBMap software package (v39.06, Bushnell 2014) with the options *ktrim=r*, *k=21*, *mink=11*, *hdist=2*, *tpe*, *tbo*, *trimq=10* and *minlen=100*. The trimmed reads were aligned to the *B. hordei* reference genome TUM1 (Liu et al. 2026) employing bwa-mem2 (v2.2.1, Vasimuddin et al. 2019). The alignments were sorted using Samtools v1.17 (Danecek et al. 2021). New read-group and library information was added to the alignment files to fulfil GATK (v.4.4.0, McKenna et al. 2010) requirements, with which the following steps were conducted. Duplicate reads were marked using GATK MarkDuplicates, and individual Genomic Variant Call Format (gvcf) files were produced with GATK HaplotypeCaller with options *–ploidy 1* and *–ERC GVCF* for each isolate and for all sites. Using GATK GenomicsDBImport a database with the variant calls for all isolates was created and a joint genotype calling was conducted from this database, using GATK GenotypeGVCFs, which resulted in a joint VCF file of all 53 isolates. Mating types were assigned by aligning the trimmed reads for each isolate to the old reference genome DH14 (Frantzeskakis et al. 2018), which carries mating type gene MAT-1-2-1. All isolates that lacked the MAT-1-2-1 region were assigned mating type MAT-1-1-1. A gene region VCF was subset from the full genome VCF file, using bedtools (v2.31.1) software and a gene region bed file as input, which was generated from the TUM1 reference annotation (v1.3.7).

For 13 isolates, ONT long-reads were obtained. These reads were aligned to the TUM1 reference genome using minimap2 (v2.28-r1209, Li 2018). Variants were called using the bcftools software suite (v1.19, Li 2011). In a first step, a multiple alignment was created using bcftools mpileup with the options *—config ont* on the aligned bam files. Then, variants were called using bcftools call with prior *-P 0.1*. The variants were filtered with bcftools filter and options *-s LowQual* and *-e ‘QUAL <* 20 || *DP >* 100*’*.

For polarizing the SNPs, isolate CHE 96224 from *Blumeria graminis* f.sp. *tritici* (*Bgt*, Kunz et al. 2023) was used as the outgroup species. *Bgt* short-read data (SRR7642212) was treated similarly to the *B. hordei* reads and variants were jointly called from a genotype database containing *Bgt* and 43 *B. hordei* isolates that were identified as coming from the same population (see below). Using a custom made Python script, sites were annotated in the VCF file as ancestral sites if reference and *Bgt* carried the same allele. The SNPs called from long-read data for 13 isolates were polarized in the same way.

### 2.3 Population Genomic Statistics

The population genomic structure was analyzed using principal component analysis (PCA). For this, the gene region VCF was converted to bed format using PLINK (v2.00, Chang et al. 2015) with options *—freq*, *—make-bed*, *—allow-extra-chr*, *—max-alleles 2* and *—set-missing-var-ids @:#*. PCA was performed with PLINK on the created bed file. The first two PCs, explaining 6.3 % and 5.9 % of the variation, were used to identify clonal isolates (5 isolates) and two possible sub- or outlier populations (3 and 2 isolates, respectively). The remaining 43 isolates were assessed to form a well-defined population, here called the core population, and were used for the subsequent analyses.

The population genomic summary statistics: nucleotide diversity (*π*), Watterson’s estimator (*θ*), and Tajima’s D were calculated for the core population using the program Pixy (v2.0.0.beta14, Bailey, Stevison, and Samuk 2025) in windows of 10 Mb along the chromosomes and also for all gene regions. Non-synonymous (*π_N_*) and synonymous (*π_S_*) sites were determined with the program SnpEff (v5.4c Cingolani et al. 2012) for the TUM1 reference and the gene annotation (Liu et al. 2026). These were used to calculate the ratio of substitution rates at non-synonymous and synonymous sites (*π_N_ /π_S_*) in the population.

LD decay was computed using PLINK (v1.9.0-b.7.11) along individual chromosomes, using options: *—r2 gz*, *—ld-window-kb 1000* and *—ld-window-r2 0.2*.

### 2.4 Polarized Site Frequency Spectrum (SFS)

The SFS of polarized SNPs, using *Bgt* as the outgroup, was calculated by a custom-made Python script which employed the packages *scikit-allel*, *pysam* and *numpy*, using different sets of sites: all sites, sites in gene regions, synonymous and non-synonymous sites. A U-shaped SFS, found on all sites, may be a signature of Multiple Merger Coalescent (MMC).

### 2.5 Demography: Sequential Markovian Coalescent

The population demography was inferred by SMC, employing the MSMC2 software (v2.1.4, Schiffels and Wang 2020). Multihetsep (mhs) files for MSMC2 input were prepared for each chromosome from the polarized VCF files. For selected chromosomes, pairs of two individuals were processed together into fake-diploid VCFs using a custom-made Python script. Sets of 21 fake-diploid chromosome VCFs were reformatted together to mhs format, using the *generate multihetsep.py* script (https://github.com/stschiff/msmc-tools) with a positive mappability mask and a negative mask excluding the repetitive regions of the genome (approx 75 %), generated from RepeatMaster output from TUM1 (Liu et al. 2026). MSMC2 was run with the options *-r 0.014* and *-p 1*2+15*1+1*2*, with rho (-r) calculated as recombination rate / mutation rate. Both values are taken from sister species *Bgt* as recombination rate = 2.8 × 10^−7^ (Müller et al. 2019) and mutation rate = 4 × 10^−7^ (Sotiropoulos et al. 2022). For the calculation of rho, the recombination rate was further divided by 5. This assumes that in the life cycle of *B. hordei* one sexual phase is followed by four asexual phases, which is similar to the estimate of 5.2 for *Bgt* (Sotiropoulos et al. 2022).

### 2.6 Neutral Demographic Simulations

Summary statistics, which could be leveraged for model choice and parameter inference, were simulated with MSPrime (Baumdicker et al. 2021). Three separate 1 Mb chromosome segments were simulated to represent subtelomeric regions of chromosomes 1, 2, and 6 of *B. hordei.* These regions were used due to the similarity of the estimated recombination rates in *Bgt* (Müller et al. 2019). A molecular recombination rate of 3.37 × 10^−7^ was used and a mutation rate of 5 × 10^−7^. A total of 195 summary statistics were computed per simulation (for the RF-ABC and NPE): the unfolded site frequency spectrum (SFS) bins 1–42, SFS quantiles, Tajima’s *D*, normalised Tajima’s 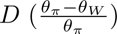, pairwise Hamming distance, linkage disequilibrium statistics (*r*^2^, interchromsomal *r*^2^, and *r*^2^_max_ equivalents (VanLiere and Rosenberg 2008)), LD frequency spectra stratified by derived allele frequency class, and adjacent-site *r*^2^. A complete list is provided in the Appendix. The addition here of LD measures between chromosomes is intended specifically to capture the potential for infinite-range LD under MMC scenarios and lower values of *α* (Birkner, Blath, and Eldon 2013; Korfmann et al. 2024b).

#### 2.6.1 Approximate Bayesian Computation (ABC)

Random Forest ABC (RF-ABC, Pudlo et al. 2015) was used to make two separate model-choice decisions regarding the detection of a *β*-coalescent model. As the results of the MSMC2 suggested a severe recent bottleneck followed by population expansion, the models of Kingman coalescence with demographic bottleneck, *β*-coalescent with constant demography, and *β*-coalescent with bottleneck were used to conduct neutral simulations to select an appropriate model for coalescence combined with demography with *α* ∼ U(1.3, 1.8). Each model was simulated 30,000 times, yielding a total dataset of 90,000 simulations.

Another model selection was performed to infer an approximate estimate for the *α* parameter of the *β*-coalescent. Five models corresponding to *α* ∈ {1.9, 1.7, 1.5, 1.3, 1.1} were used to generate neutral simulations with a growth parameter ranging between expanding and contracting demographies with a *β*-coalescent process. *α* = 1.9 was used to represent the *β*-coalescent as it approaches a Kingman process. For each *α*-value we generated 25,000 simulations for a total of 125,000 simulations.

#### 2.6.2 Neural Posterior Estimation (NPE)

A NPE (Papamakarios and Murray 2016) from the Simulation Based Inference Python package sbi (Tejero-Cantero et al. 2020) was used to jointly infer coalescent and demographic parameters under the *β*-coalescent model. A total of 100,000 neutral simulations were generated using the same genomic configuration described above (three 1 Mb segments; *µ* = 5×10^−7^; *ρ* = 3.37×10^−7^; *n* = 43 haplotypes), with parameters drawn from a uniform prior: *α* ∼ U(1.3, 1.9), a scaling nuisance parameter *k* ∼ U(1, 150) representing the ratio of asexual to sexual generations, which scales the effective recombination rate as *ρ*_eff_ = *ρ/*(*k* + 1) to absorb uncertainty in the relative frequency of sexual reproduction and molecular rates, and relative effective population sizes *N_e,i_* ∼ U(0.1, 5) for 11 discrete time windows with breakpoints at 10, 50, 100, 200, 300, 400, 500, 1,000, 10,000, 100,000, and 1,000,000 generations before the present. The *N_e,i_* values are expressed as multiples of the present-day *N_e_*, representing a demographic trend rather than absolute census sizes. Simulations with fewer than 2,860 or more than 2,900 segregating sites were rejected to match the observed data. The conditional density estimator was implemented as a Neural Spline Flow (NSF) with 64 hidden features and five coupling transforms, trained via NPE.

#### 2.6.3 Posterior calibration via Simulation-Based Calibration

We assessed posterior calibration using Simulation-Based Calibration (SBC; Talts et al. 2020). SBC exploits a self-consistency property of the Bayesian joint distribution: if the trained density estimator *q_ϕ_*(***θ*** | ***x***) recovers the true posterior, then for parameter values drawn from the prior, the rank of each true parameter among posterior samples drawn from its corresponding inferred posterior is uniformly distributed on {0, 1*, …, L*}, where *L* is the number of posterior samples per trial. Departures from uniformity diagnose specific failure modes: U-shaped rank histograms indicate posteriors that are too narrow (overconfidence), inverted-U shapes indicate posteriors that are too wide (underconfidence), and monotonic slopes indicate systematic bias.

For each of *N* = 500 SBC trials we (i) drew a parameter vector ***θ***^(*i*)^ from the prior, (ii) simulated a synthetic dataset ***x***^(*i*)^ using the same msprime configuration as the training simulations, (iii) drew *L* = 1,000 samples from *q_ϕ_*(***θ*** | ***x***^(*i*)^), and (iv) computed, for each parameter dimension separately, the rank of ***θ***^(*i*)^ among the posterior samples. Uniformity of the resulting rank histograms was assessed visually and indicates well-calibrated marginal posteriors. We note that uniform ranks are also produced when the marginal posterior is essentially the prior (when the data are uninformative for that parameter), since the true value then falls at a random rank by construction. SBC therefore certifies that the reported uncertainty is honest, but it does not by itself establish identifiability; we address the latter via posterior-width comparisons in the results.

#### 2.6.4 Posterior accuracy and confounding diagnostics

Beyond calibration, we assessed how accurately the NPE recovered the true *α* used to simulate each SBC trial by computing the Pearson correlation between the true values 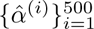 and the posterior means 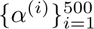. This correlation is a direct measure of point-estimate accuracy across the full prior range and is a strictly stronger condition than calibration: a posterior can be calibrated yet uninformative, in which case posterior means will not track true values and the correlation will be near zero.

To test whether the inferred *α* was systematically confounded by demographic history, we computed the Pearson correlation between each true window-specific population size 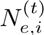 used to generate trial *t* and the corresponding posterior mean â^(*t*)^, across all 500 trials and all 11 time windows. A nonzero correlation would indicate that inferred *α* depends on a simulation’s demographic history independently of its true *α*, *i.e.* that demographic and coalescent signals are entangled in the summary statistics used by the NPE. Significance was assessed by permutation, using 1,000 random permutations of the true *N_e,i_*values within each time window.

### 2.7 Selective Sweep detection

Detection of selective sweeps was conducted with two different software suites, SweeD (v4.0.0, Pavlidis et al. 2013) and RAiSD (v2.9, Alachiotis and Pavlidis 2018). For real data, each chromosome was scanned, starting from the VCF files for RAiSD. For SweeD the input had to be reformatted to sweepFinder format using a custom-made Python script and then scanned with the option *-grid = 1000*. We first used an outlier approach, taking the top 0.1% RAiSD higher values, yielding a threshold of 246.48, and the top 1% SweeD higher values with threshold of 0.50.

For sweep validation, the simulated demography and *α* parameter that produced the best fit to the observed SFS were used to run a neutral demographic baseline and to generate a minimum cutoff value for sweep detection. A recombining sub-telomeric region and centromere region with 1% of the recombination rate of the sub-telomeric region were simulated. Sweep detection was conducted as done for the real data. No significant sweep was detected with SweeD on the simulated data. The 0.1% RAiSD significant threshold was *µ* = 7.01, and six sweep regions were identified as above this threshold in this 1Mb simulated region. We used a conservative threshold of *µ* = 8 to adjust the genome-wide significance ratio on the real data.

## 3 Results

3.1 The German *B. hordei* population

The 53 *B. hordei* isolates were collected from barley plants during the growing season (Table S1 and Figure S1). We expect isolates to be part of the same population, given previous results about the large dispersal ability of *Blumeria* species (Dreiseitl 2019; Jigisha et al. 2025). Isolates were purified in two rounds of propagation and then sequenced using short-read technology for all samples. Moreover, long-read sequencing technology was employed for a selected subset of 13 isolates. From the short-read data, aligned to the near-complete reference genome TUM1 (Liu et al. 2026), 679,749 variant sites were detected, of which 647,713 were bi-allelic SNPs. A PCA analysis of the bi-allelic SNPs in the gene regions (74,664 SNPs) revealed very little population structure and that the majority of isolates form a main cluster (Figure 1, magnification on the right). Isolates falling outside the cluster (five isolates) or clones of other isolates (judged by their near-identical positions in the PCA; five isolates) were removed from the data set. The mating type distribution was quite balanced among the collected accessions. Of the 48 non-clonal isolates, 26 were of mating type MAT1-1-1 and 22 of mating type MAT-1-2-1, or for the core population 23 and 20, respectively.

**Figure 1:**
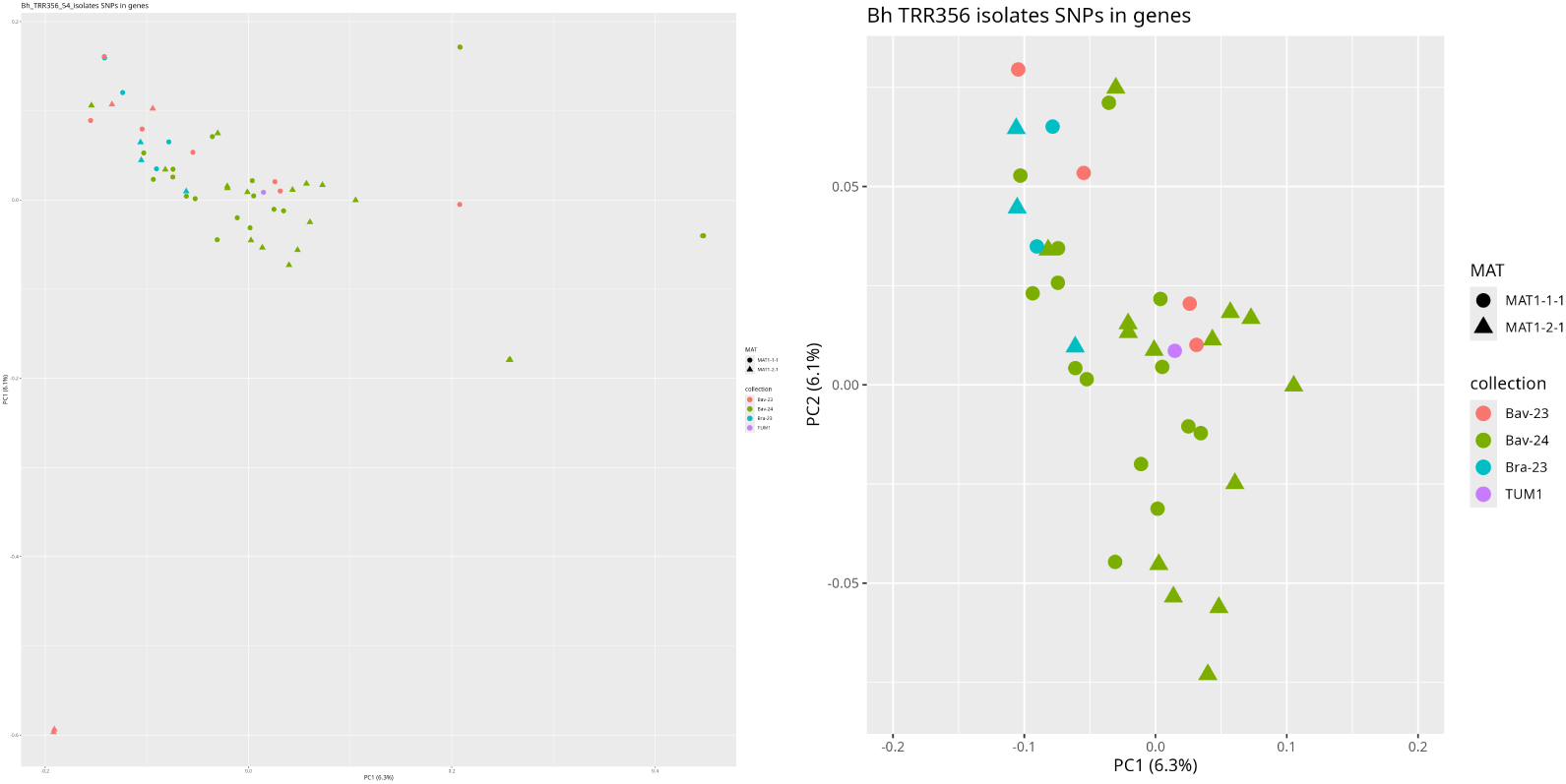
PCA using gene based SNPs from short-read genomic data. Plot of PC1 vs. PC2, Left: all isolates, right: magnification of the main cluster. Colour of points indicate the federal state and year of sampling. TUM1 is the reference isolate. The shape of the points reflects the mating type (circle: MAT 1-2-1, triangle: MAT 1-2-2).

The following analyses were conducted using the dataset for the 43 isolates of the main cluster (Table S1).

### 3.2 Site Frequency Spectrum (SFS) and past demography

To gain insight into the population dynamics, the polarized SFS of the gene regions (*Bgt* as outgroup) was determined both using the short-read data set and additionally a similar generated data set, using long-read data for 13 *B. hordei* isolates (Figure 2). The use of long-read data seemed advisable as the *B. hordei* genome carries a very high content of transposable elements (TEs). Repeat detection software identifies over 70 % of the genome as repeat regions (Frantzeskakis et al. 2018; Liu et al. 2026). Several transposable elements are present in very large numbers, which could lead to errors in the allele frequency statistics deduced from short-read data, as reads are only 150 bp long and may be misaligned in repeat regions. In contrast, alignments of long sequencing reads have less uncertainty even in repeat regions, and the resulting allele frequencies is expected to be more accurate. With both types of sequencing reads, a U-shaped SFS was found. The U-shaped was also observed when analyzing full genome, only synonymous or only non-synonymous SNPs (Figure S8). One possible evolutionary process that can lead to a U-shaped SFS is strong positive pervasive selection (at many targets in the genome). To follow this possibility, Tajima’s D was determined (Figure S4), which was on average only slightly negative, and perhaps not as strongly negative, as could be expected if only strong positive selection was at work.

**Figure 2:**
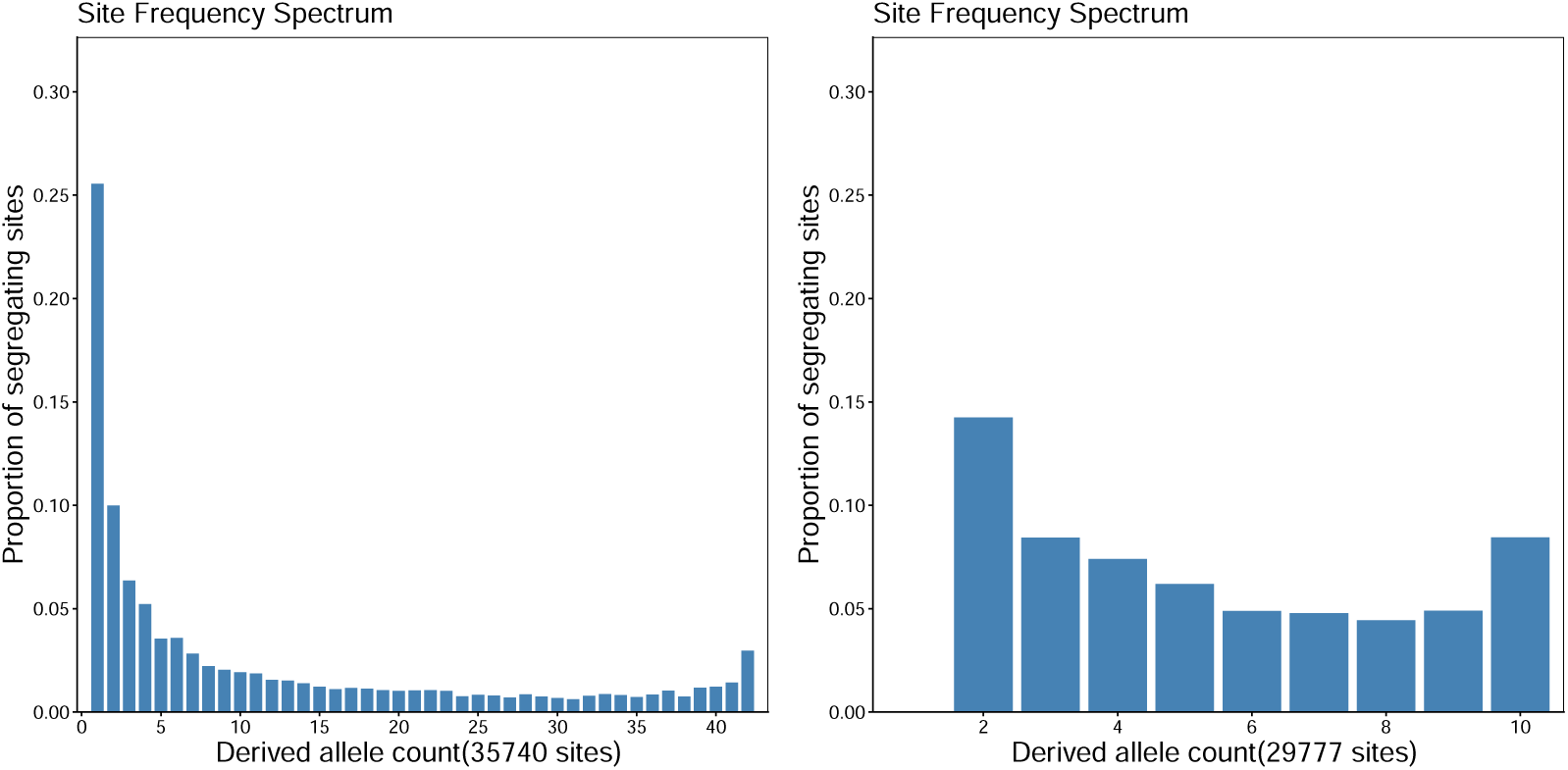
Folded SFS of the gene regions of the *Bh* population using short read data from 43 isolates (left) or long-read data from 11 isolates (right).

Another process leading to a U-shaped SFS could be a recent bottleneck, which should be detectable when inferring the population’s past demography. The demography was based on SNPs from non-repetitive regions, and repeated for several chromosomes (Figure 3). The inferred demographies of the chromosomes were quite similar and revealed a recent population extension in the last ca. 70 years. Before that, a population contraction in the last 1,000 up to 2,000 years was revealed. Going further back in time, the *B. hordei* population seemed not to have undergone large population changes. We caution not to interpret the results from more than 10^5^ years ago, as the coalescent methods carry a greater uncertainty at extreme ends of the time / generation scale.

**Figure 3:**
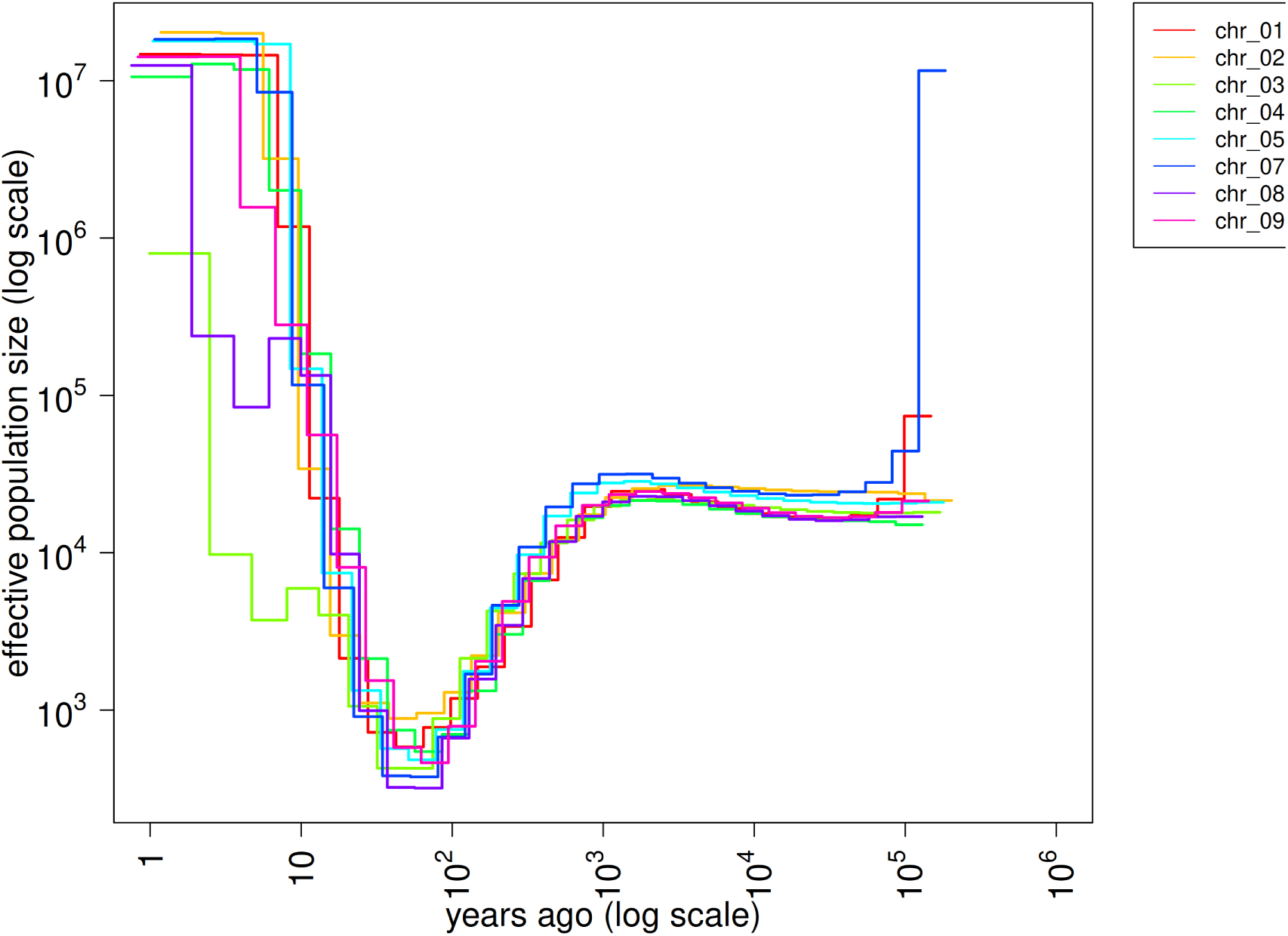
Inferred demography of the population using MSCM2 on short-read polymorphism data from eight different chromosomes.

As population structure was already ruled out as a possible other reason for a U-shaped SFS, the most likely explanation was a MMC, either due to neutral or selective sweepstakes reproduction.

### 3.3 Reliance on SFS signal favors Kingman coalescence

The polymorphism pattern most often associated with MMC is a U-shaped SFS with an excess of rare (singleton) alleles (Schweinsberg 2003; Tellier and Lemaire 2014; Eldon et al. 2015; Menardo, Gagneux, and Freund 2021; Freund et al. 2023). While an excess of rare alleles can be a consequence of rapid population expansion or MMC, leveraging this signal with information in the right tail of the SFS (the presence of a U-shape) has been shown to allow for the differentiation of expansion or multiple-merger models by itself or combined with other summary statistics (Koskela 2018; Koskela and Wilke Berenguer 2019; Freund and Siri-Jégousse 2021).

To determine if *B. hordei* is best described by a Kingman or *β*-coalescent model, a model choice was conducted with random forest ABC (RF-ABC, Pudlo et al. 2015). The simulations were performed to account for the uncertainty of the demographic history of *B. hordei*, including a random uniform growth parameter, which could simulate a range of demographic histories from population contraction to expansion. The model choice was performed for the *α* parameter of the *β*-coalescent for five different models of *α* ∈ {1.9, 1.7, 1.5, 1.3, 1.1}.

The RF-ABC trained with summary statistics, which include various information from the SFS, showed a strong ability to differentiate *α* = 1.9 from the rest of the multiple merger models with an Out-Of-Bag error of 5.17% (Figure 4). When assigning the observed data to one of the models, the posterior probability is 62.6% for *α* = 1.9, suggesting a Kingman coalescent process (Figure 10). However, when looking at the most important variables for training the random forests, a strong dependency on the singleton class (SFS1) is seen, as well as various representations of the SFS such as the SFS quantiles and Tajima’s D. This suggests a potential oversensitivity of the RF-ABC to singletons in particular, but also representations of the SFS more generally (Figure 11).

**Figure 4:**
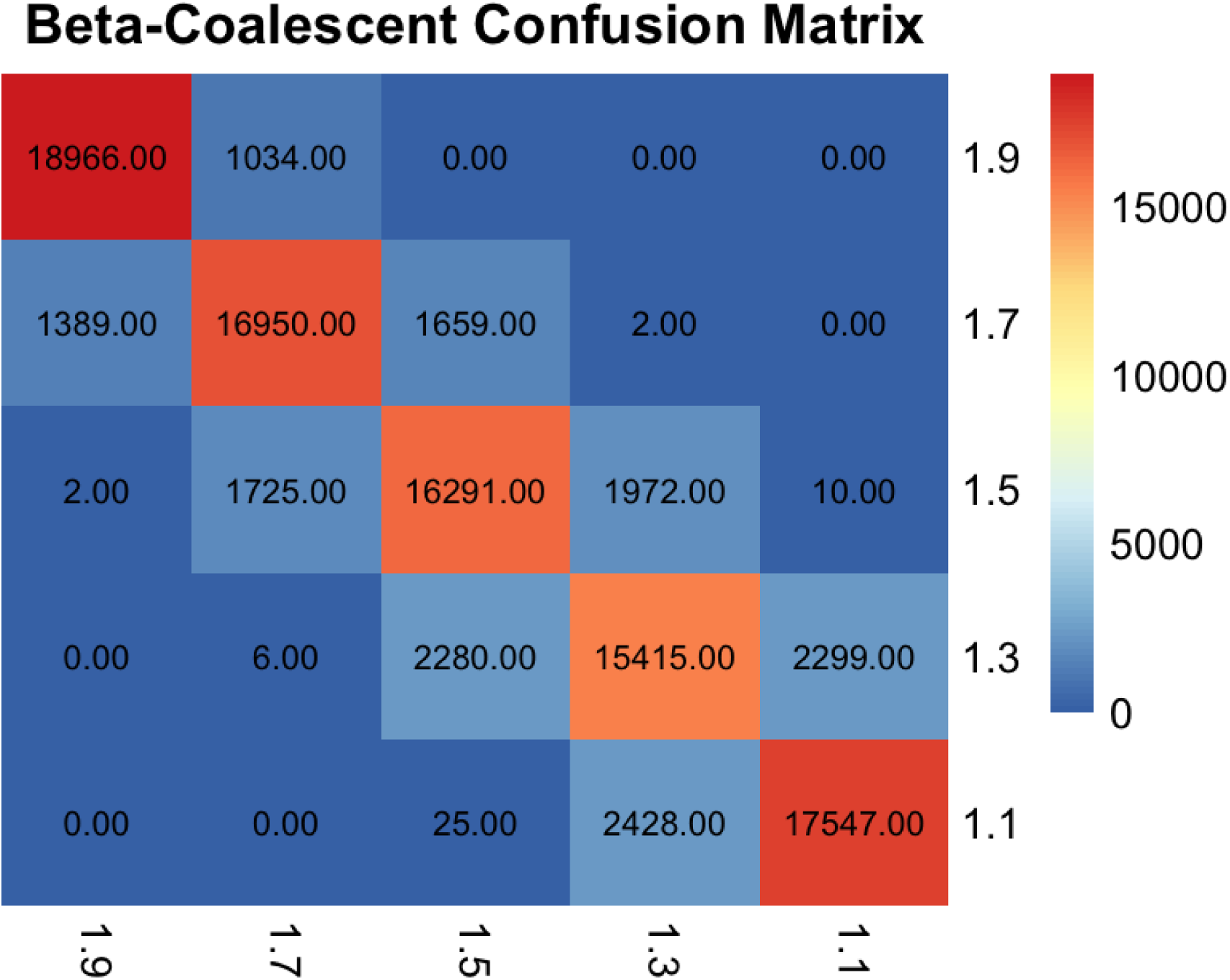
Confusion matrix of random forests for models of different *α* parameters. Out of bag errors: 5.17% for *α* = 1.9; 15.25% for *α* = 1.7; 18.55% for *α* = 1.5; 22.93% for *α* = 1.3; 12.27% for *α* = 1.1.

### 3.4 Kingman-based demography and artifacts caused by MMC

A known effect of the MMC is that mis-specification and use of demographic inference methods with assumptions of a Kingman coalescence process, can result in the inference of population expansion (Menardo, Gagneux, and Freund 2021; Korfmann et al. 2024b). This is an artifact caused by the star-shaped tree topologies, which are caused by population expansion under the Kingman coalescent, being confounded with the similar topologies caused by a MMC.

Star-shaped genealogies have the effect of increasing the prevalence of rare alleles. However, as seen in Figure 9, the SFS from the regions used for the ABC exhibited a low proportion of rare alleles with singletons representing only 27% of total SNPs, and a mean Tajima’s D of −0.34. If the frequency of rare alleles would be an accurate indicator of Kingman coalescence, a Kingman-based demographic inference should display a demographic history that is not especially characterized by star-shaped genealogies based on the lack of rare alleles in our observed sequence.

While many demographic inference methods rely on the SFS, one would expect that if the underlying model of the observed sequence is truly a Kingman coalescent, then inference methods that assume a Kingman coalescent, but rely on polymorphism signals other than the SFS, should yield a demographic history concordant with interpretations of the SFS. MSMC2 was used to fit a demographic model based on the density of pairwise SNPs along the genome, and thus should be able to detect star-shaped genealogies with a polymorphism signal independent of the SFS characteristics.

When using the MSMC2, the inferred demographic history was reflective of a star-shaped genealogy with an extreme recent bottleneck and population expansion (Figure 3). Such an extreme demography, where over 99% of lineages enter a bottleneck, was discordant with the observed sequence having relatively few rare alleles and very moderately negative Tajima’s D. When performing a similar model choice RF-ABC between a Kingman coalescent with bottleneck informed by the MSMC2 results, *β*-coalescent with a flat demography, and a *β*-coalescent with bottleneck, the *β*-coalescent models received 85.6% of the model votes, with the selected model being a constant demography *β*-coalescent with 47.4% of the model votes (Figure 16). This suggested that an extremely skewed Kingman-based demography does not adequately describe the data.

### 3.5 SFS-based statistics are biased in the joint presence of MMC and demographic events

While most methods for detecting MMC rely on SFS information, the confounding effects of demographic history combined with a multiple-merger process on the SFS have not been explored until now. To illustrate this, we ran 20 simulations for four different conditions of recent demographic changes combined with MMC, and then plotted the corresponding SFS. The effect of recent demographic history on the SFS was pronounced, calling into question the ability of SFS-based methods to accurately infer MMC in the presence of changing demographic histories (Figure 7).

To explore the robustness of the RF-ABC against this potential confounding effect, the model choice was run four additional times with incrementally fewer statistics representative of the SFS each time: 1) no singletons, 2) no SFS bins, 3) no SFS bins or quantiles, 4) no SFS information or Tajima’s D. While the overall Out-Of-Bag (OOB) error for each RF-ABC increased as SFS information was removed, the ability to distinguish between the Kingman and the multiple-merger approximation remained stable (∼5.18–5.5% classification error). As SFS information is removed, all predictions for each of the additional RF models supported a moderate multiple-merger process of *α* = 1.7 (Figure 14).

Given the apparent sensitivity of the SFS to demographic changes and the risk of SFS information causing over-fitting of the random forests (in RF-ABC), we question how much bias is being introduced by current inference methods, which leverage almost exclusively the signals of the SFS. A simple parameter inference for demographic history and MMC using the RF-ABC would also be of limited efficacy. Indeed, we suggest that RF-ABC is trained for each inferred parameter individually, so that parameters for *α* and demographic changes would be potentially confounded. As such, a framework for simultaneous parameter inference would be best suited for disentangling the effects of MMC and demographic history.

### 3.6 Neural Posterior Estimation of *β*-oalescent illustrates potential demographic complexity

Given the limitations of the RF-ABC highlighted above and the inability to infer joint posterior probabilities for MMC and demographic parameters, a Neural Posterior Estimation (NPE) was constructed (Papamakarios and Murray 2016). NPE trains a conditional density estimator *q_ϕ_*(***θ*** | ***x***), parametrized by a neural network with weights *ϕ*, to approximate a posterior directly. It offers the benefit of joint parameter inference, which is advantageous given the hypothesis of confounding effects for *α* and demographic history. In comparison to the RF-ABC, the NPE can also handle much larger summary statistic sets without the need to prune uninformative statistics or potentially collinear predictors.

A larger parameter space was also employed than in the previous RF-ABC, including *α* ∼ U(1.3, 1.9). A nuisance parameter *k* represented the ratio of asexual to sexual generations by scaling the effective recombination rate, and we modeled a relative effective population size (*N_e_*) for 11 discrete time windows from 10 to 10 million generations ago. As the *α* parameter affects the population scaling, these *N_e_* values are represented as a multiple from 0.1 to 5 times the present day *N_e_*, thus sketching a historical demographic trend, rather than a purely numerical *N_e_*estimate. A Neural Spline Flow (NSF) architecture was used as the density estimator, which is capable of representing multimodal and asymmetric posteriors that a standard Masked Autoregressive Flow (MAF) would over-smooth.

Posterior calibration of the trained NSF was assessed using Simulation-Based Calibration (see methods). The rank histograms for *α* and the nuisance scaling parameter *k* were consistent with uniformity (Figure 12), indicating well-calibrated marginal posteriors for the parameters of primary interest. The 11 time-windowed *N_e_*parameters also produced approximately uniform rank histograms. SBC alone could not distinguish two explanations for this pattern: (i) the *N_e_*posteriors were calibrated and informative, or (ii) the *N_e_* posteriors tracked the prior closely, in which case the true value fell at a random rank by construction. The marginal *N_e_* posteriors (Figure 15) had credible intervals that did not substantially narrow relative to the prior, identifying the second scenario as the operative one. We interpret this as the model correctly acknowledging its uncertainty over historical population sizes given the available summary statistics, rather than producing falsely confident *N_e_* estimates. The *N_e_* windows functioned as nuisance parameters that absorb demographic complexity without being individually identifiable.

Accounting for demographic complexity, we found evidence for MMC in *B. hordei* (*α* = 1.627 ± 0.116, median 1.662, 95% CI [1.379, 1.837]), suggesting a moderate multiple-merger process clearly distinct from the conservative Kingman approximation (*α* = 1.9). The scaling parameter *k* had mean 3.608 ± 1.949 (median 3.179, 95% CI [1.089, 7.706]), suggesting a reasonably selected recombination and mutation rate given the uncertainty in the ratio of asexual to sexual generations. These estimates were consistent with those obtained from a MAF density estimator (*α* = 1.645, 95% CI [1.42, 1.86]), supporting the robustness of the *α* inference to the choice of neural architecture.

The NSF posterior nonetheless revealed pronounced multimodality in the *N_e_* marginals that the MAF approximated as smooth unimodal distributions, reflecting potential degeneracy: multiple combinations of *α* and population-size trajectories are consistent with the observed summary statistics. Despite this multimodality, the 95% credible interval for *α* excludes the conservative Kingman approximation *α* = 1.9, so our NPE framework confidently infers MMC even after marginalizing over the demographic nuisance parameters.

NPE accuracy across the full prior range was assessed using the same 500 SBC trials. The Pearson correlation between the true *α* values used to generate each trial and the corresponding posterior mean *α*^ was *r* = 0.81 (*p <* 0.001). This indicated that the NPE recovered *α* accurately across the full U(1.3, 1.9) prior range and across the wide range of demographic histories drawn during the SBC procedure. Calibration and accuracy are complementary diagnostics: calibration verifies that the reported uncertainty is honest, while the correlation verifies that the posterior mean is informative. The NSF posterior density estimation satisfied both criteria for *α*.

To test whether the inferred *α* was confounded by the underlying demographic history of each simulation, we computed the Pearson correlation between each true *N* ^(*t*)^ window value and the posterior mean *α*^^(*t*)^ across the 500 SBC trials. The intermediate-timescale windows *N_e,_*_2_, *N_e,_*_3_, and *N_e,_*_4_ (50–300 generations ago) showed small but statistically significant positive correlations with inferred *α* (*r* = 0.19, 0.15, and 0.13 respectively; all *p <* 0.01 by 1,000-permutation test). That is, when a simulation was generated with a larger population size at intermediate timescales, the NPE tended to infer a slightly more Kingman-like *α*, and bottlenecks at intermediate timescales coincided with lower inferred *α*. This was consistent with the SFS-based confounding mechanism: elevated *N_e_* between 50 and 300 generations ago raises the proportion of intermediate-frequency alleles, partially mimicking the less U-shaped SFS expected at higher *α*. The effect is modest, however, with these correlations jointly explaining less than 4% of the variance in inferred *α*. All other *N_e_* windows, including the most recent (*N_e,_*_1_; 10–50 generations) and all ancient windows (*N_e,_*_8_ through *N_e,_*_11_; *>* 1,000 generations), were uncorrelated with inferred *α* (|*r*| *<* 0.05, all *p >* 0.1; Figure 13).

### 3.7 Simulated SFS displays confounding effects

While the multimodality and low confidence of the *N_e_*posterior distributions showed the potential complexity of the demographic history in *B. hordei*, simulations conducted in such a highly complex parameter space are still informative. By selecting the simulation which minimized the Euclidean distance between the simulated SFS and the observed SFS, we showed the clear issue with SFS-dependent inference of the *β*-coalescent, as well as revealed the demographic scenario most well fitting the data.

The simulation with the best SFS match displayed an estimate of *α* = 1.641, consistent with the posterior mean of 1.627. The demographic model producing this SFS most strikingly showed a large population size increase around 300 years ago, followed by a rapid decline around the time of the Green Revolution, coinciding with the widespread adoption of fungicides and resistance breeding (Figure 5).

**Figure 5:**
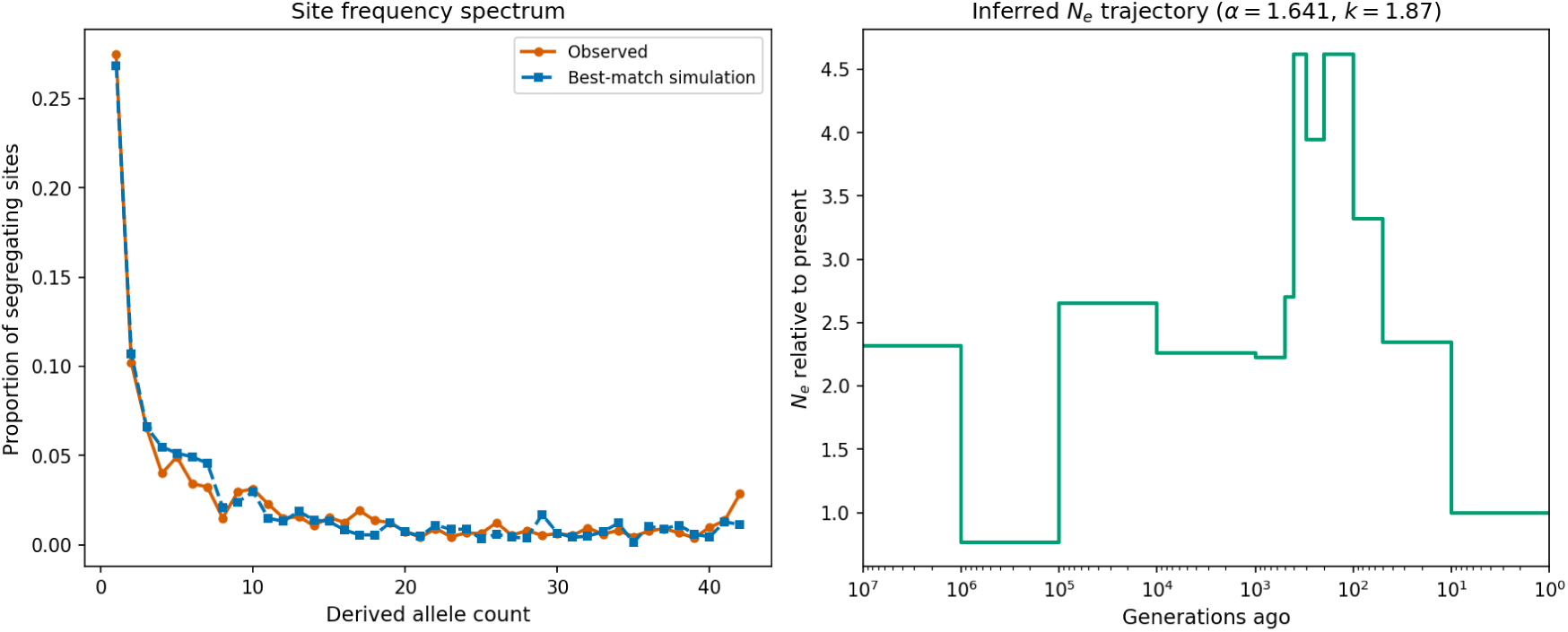
Left: The simulation’s SFS which minimizes the Euclidian distance to the observed SFS. **Right:** The simulation demographic history and *α* value which produced the closest matching SFS.

**Figure 6:**
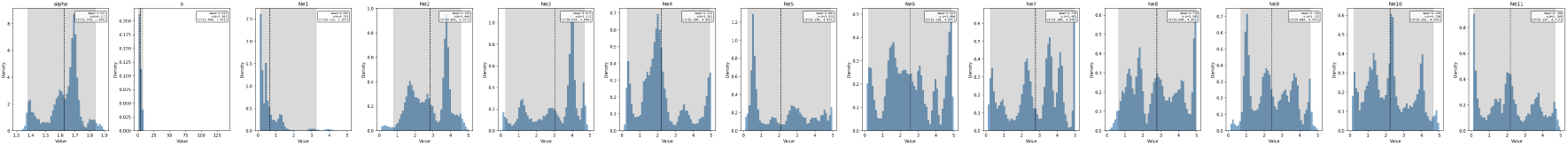
Neural spline flow posterior estimations for *α*, ***k***, and relative *N_e_*estimates for discretized time windows.

We additionally showed that the SFS from this demographic model is distorted when compared to the SFS simulated from the same *α* value under a constant demographic size. The singleton class in particular shows a 10% difference in frequency, meaning methods that rely on the SFS would be biased by inferring a much more Kingman-like coalescent process when not accounting for the demographic history (Figure 8).

### 3.8 Genome-wide scan for selective sweeps in *Blumeria hordei*

We performed genome-wide scans for signatures of selective sweeps across all 11 chromosomes of *B. hordei* using two complementary approaches: RAiSD (Raised Accuracy in Sweep Detection) and SweeD (Sweeps Detector). Both methods identified multiple candidate regions under positive selection, with some concordant and some method-specific signals The RAiSD *µ* statistics revealed pronounced selective sweep signals across the genome, with values reaching up to 1,500 in the most extreme cases. Notably, several of the highest RAiSD signals were located in pericentromeric regions (indicated by grey shading) and near the telomeres. Additional moderate-to-strong signals (RAiSD *µ >* 500) were distributed across most chromosomes, with chromosome 4 displaying multiple peaks of varying intensities, and chromosome 10 showing a significant sub-telomeric peak. Chromosomes 3, 6, and 11 showed relatively fewer strong signals, though chromosome 11 exhibited a notable peak near its telomeric region (Figure 7). Using such an outlier approach, taking the top 0.1% RAiSD higher values (threshold was 246.48), yielded approximately 1,600 outlier putative selected regions. The 1% SweeD significant threshold was 0.50 and yielded approximately 120 such regions.

**Figure 7:**
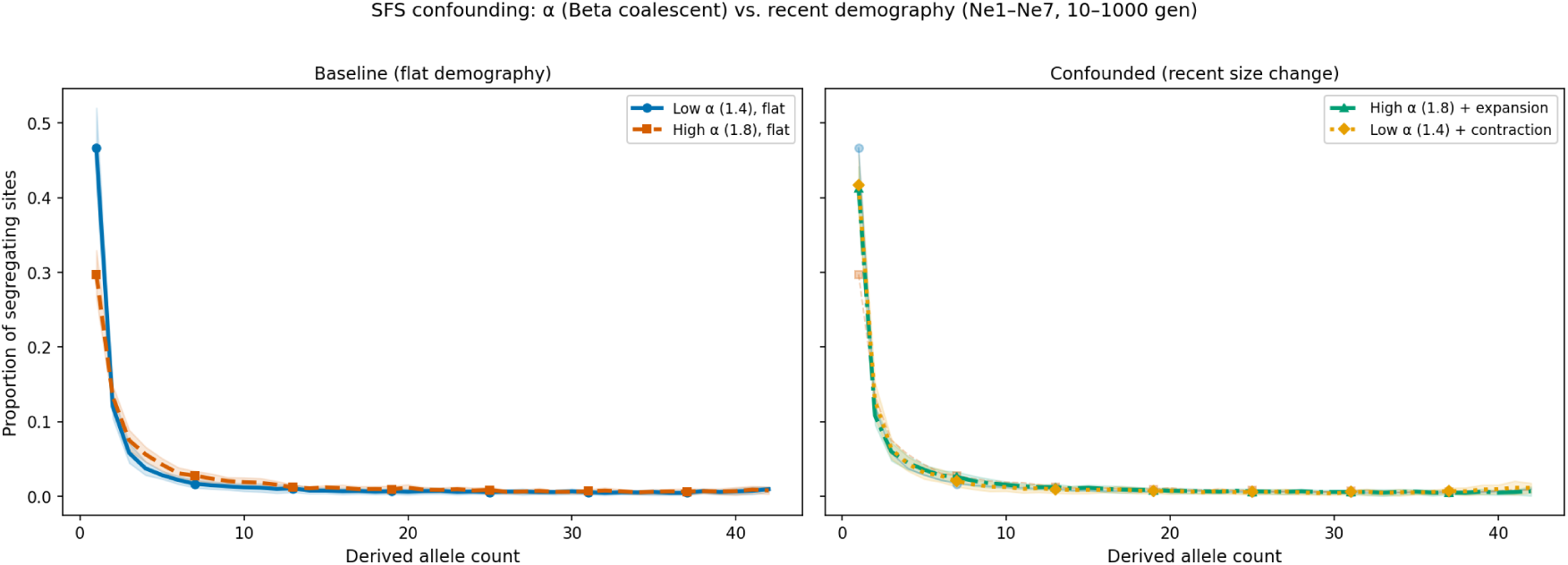
Left: Simulated SFS for constant demography at *α* = 1.4 and *α* = 1.8. **Right:** Simulated SFS for the same *α* values, but with a population size change 1000 generations ago. Baseline values are displayed to show the magnitude of SFS distortion when demography is added.

**Figure 8:**
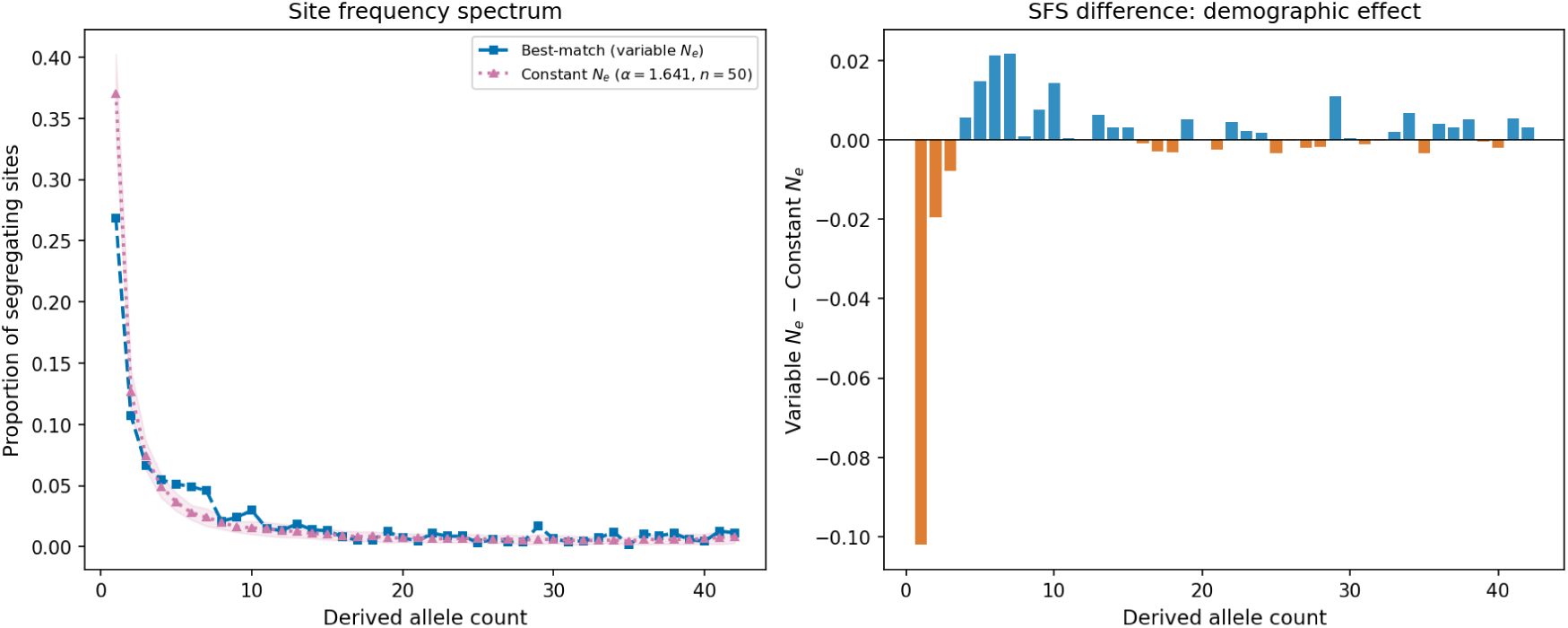
Left: The best match SFS and mean SFS for a simulation with the same *α* but constant demography. **Right:** The differences in the frequencies for each bin of the SFS when the best match SFS is substracted from the SFS generated from the same *α* value with constant demography.

Generally, centromeric and telomeric regions are masked when using RAiSD, as low-recombination regions can yield signals that mimic positive selective sweeps. However, we left them in to compare the results produced by the neutral simulation validation with the *β*-coalescent. It should be noted, though, that such large spikes in these low-recombination regions are likely artifacts, rather than genuine selective signals.

The SweeD composite likelihood ratio (CLR) analysis provided a complementary view of selective sweep signatures, with maximum CLR values reaching approximately 3.5. Notable peaks were distributed across all chromosomes, although some peaks also occurred in centromeric regions and close to the telomeres. Interestingly, while some regions showed concordance between methods (such as strong signals on chromosome 6), there were also method-specific detected regions (Figure S5).

The neutral simulations with MMC and demographic history did not yield significant signals of selection in either RAiSD or SweeD, suggesting that the methods are robust to at least moderate neutral multiple-merger processes (*α >* 1.6) with demography. Notably, the RAiSD validation showed lower *µ* values in the centromere than the sub-telomeric regions, suggesting the peaks observed in low recombination regions were an artifact with causes not necessarily related to the multiple-merger process alone, but more probably an interaction of low recombination and recurrent purifying/background selection (Figure 9). None of the RAiSD *µ* values exceeded 8.0. If this value was used as a significant threshold in the RAiSD scan of the real data, 189,597 sweep regions would be identified (11.6 % of all regions), which can be seen as the result of possible genome-wide pervasive positive selection (Figure S10.

**Figure 9:**
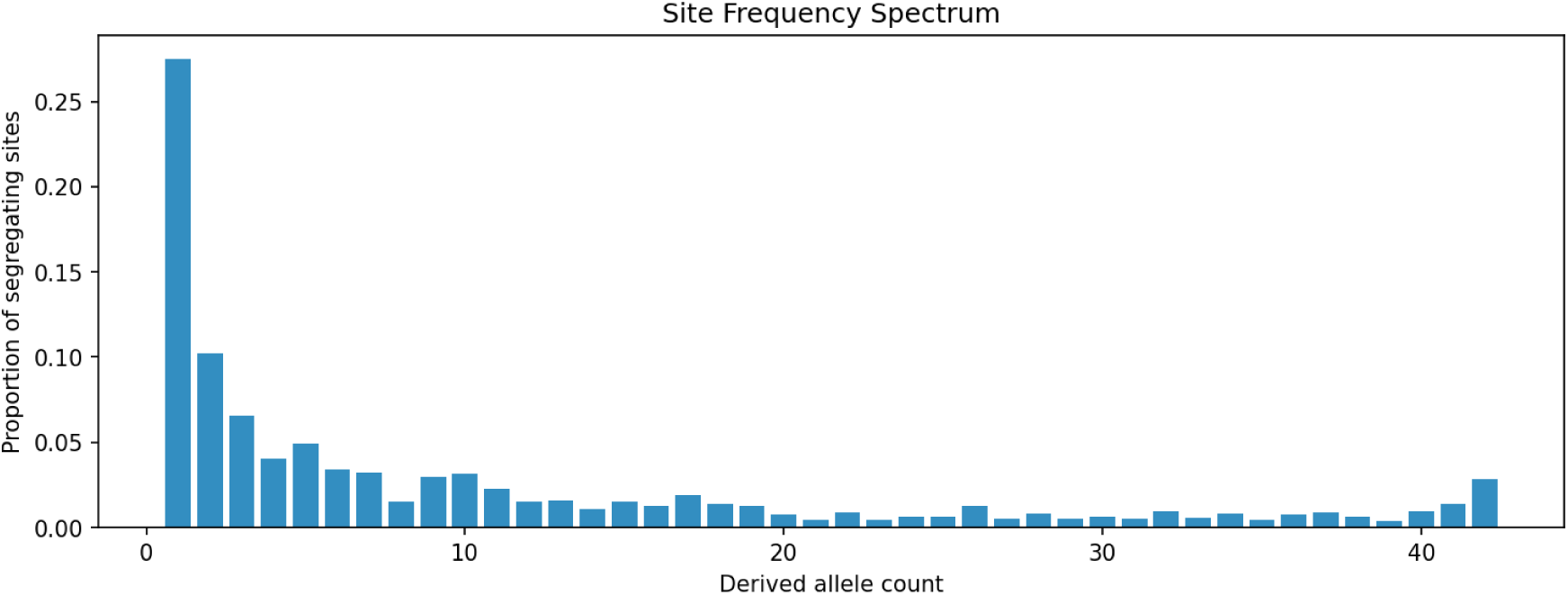
The observed SFS from sub-telomeric regions of chromosomes 1, 2, and 6.

**Figure 10:**
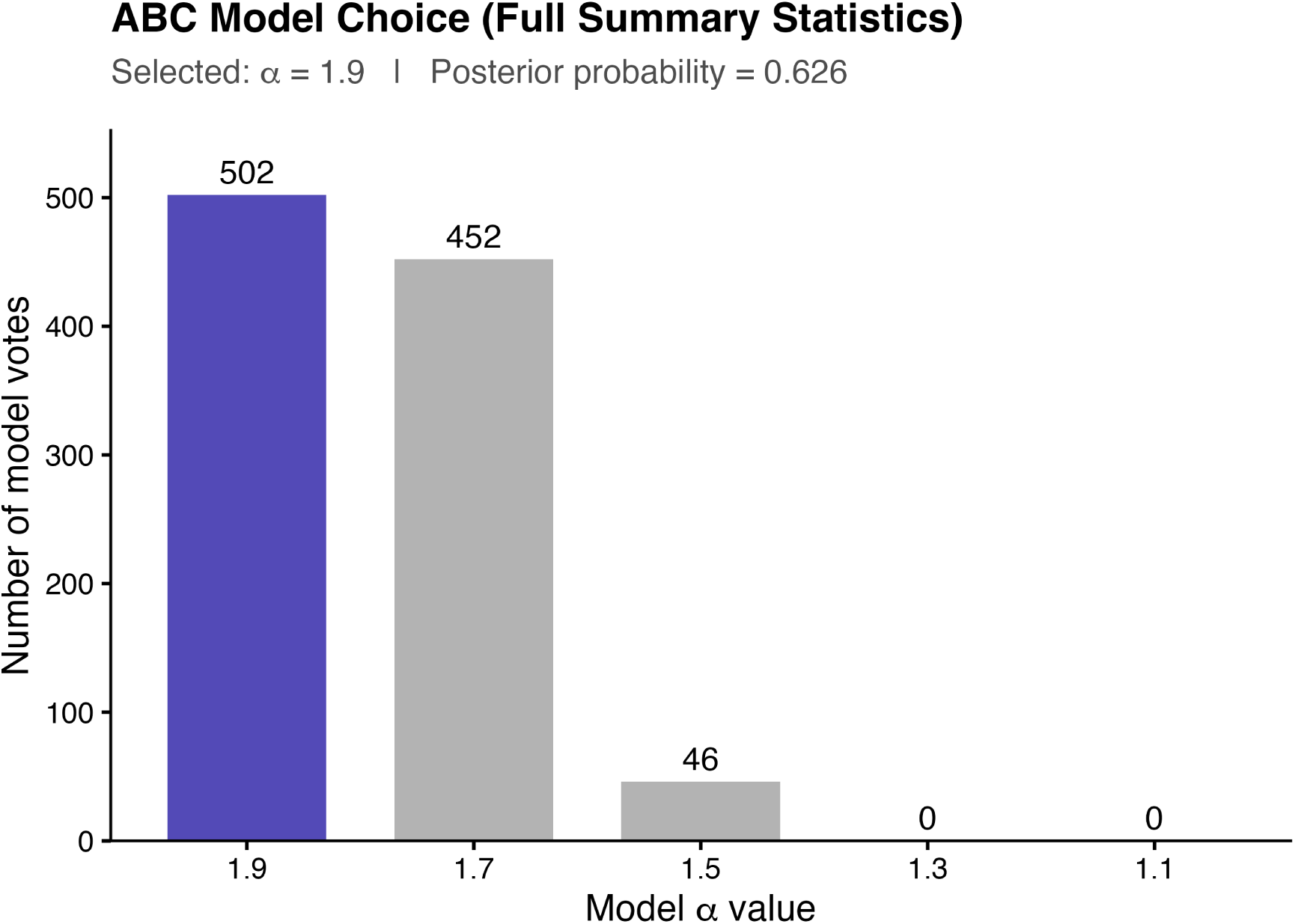
When using a set of summary statistics which includes the bins of the SFS, the RF-ABC assigns the observed data to a Kingman-like model.

**Figure 11:**
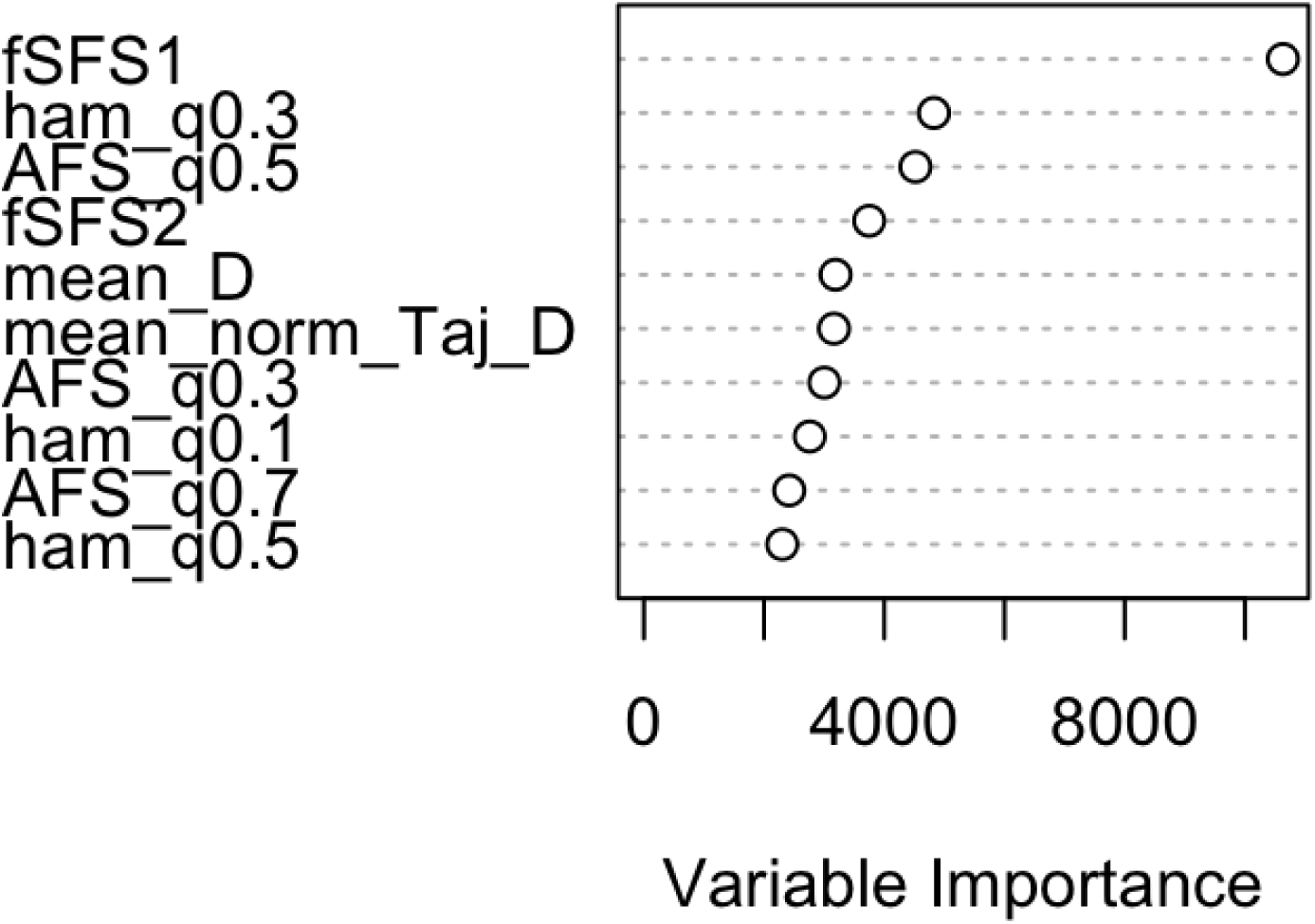
Variable importance as assigned by RF-ABC. The most important summary statistics are bins of the SFS (fSFS1 and 2), quantiles of the SFS (AFS q), mean Tajima’s D and normalized Tajima’s D, and the pairwise hamming distance quantiles (ham q).

**Figure 12:**
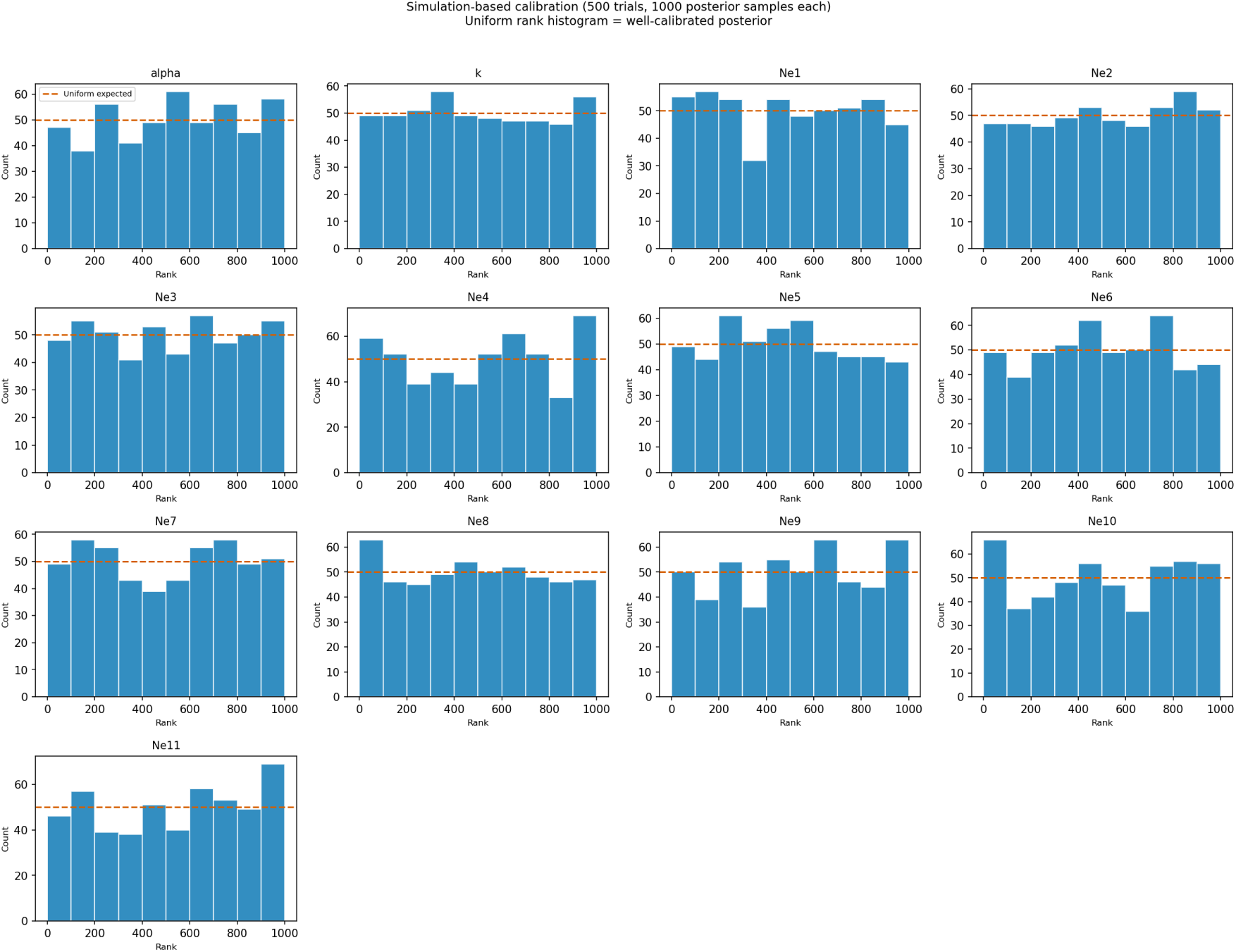
Simulation-Based Calibration (SBC) rank histograms for all 13 inferred parameters across 500 prior-spanning trials (1,000 posterior samples per trial). The dashed orange line indicates the expected count per bin under a uniform rank distribution, which is the criterion for a well-calibrated posterior. Bars above or below this line indicate over- or under-representation at those ranks. *α* and *k* show rank distributions consistent with uniformity, confirming well-calibrated posteriors. The *N_e_* window parameters also show approximately uniform distributions, reflecting calibrated but uninformative posteriors: posteriors that are nearly as wide as the prior will produce uniform rank statistics regardless of whether the parameter is being accurately inferred.

**Figure 13:**
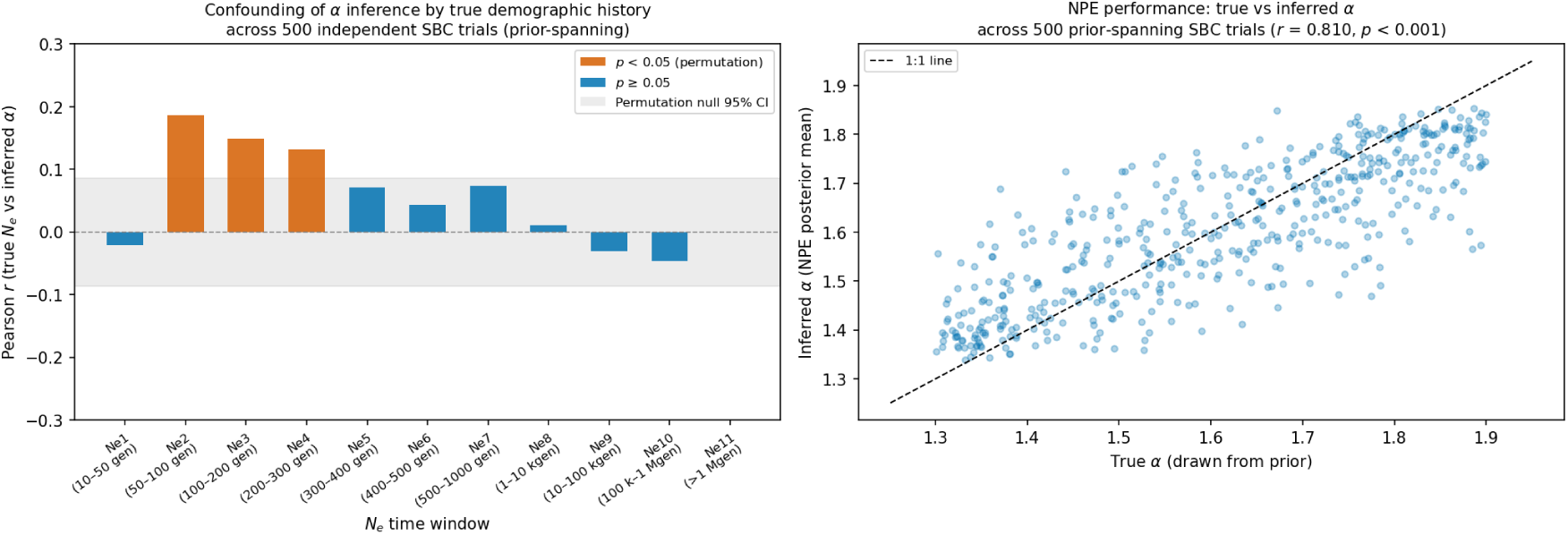
Assessment of demographic confounding in NPE *α* inference using 500 prior-spanning SBC trials. *Left:* Pearson correlation between the true *N_e,i_* value in each time window and the posterior mean *α* inferred by the NPE. Orange bars indicate windows with statistically significant correlations (*p <* 0.05, permutation test); the shaded band shows the permutation null 95% CI. *N_e,_*_2_–*N_e,_*_4_ (50–300 generations ago) show small positive correlations (*r* = 0.13–0.19), indicating that larger intermediate-timescale population sizes cause a modest upward bias in inferred *α*, consistent with elevated *N_e_* at these timescales generating intermediate-frequency alleles that mimic a more Kingman-like SFS. All other windows are uncorrelated with inferred *α*. *Right:* Pearson correlation between true and NPE-inferred posterior mean *α* across all 500 trials (*r* = 0.81, *p <* 0.001), demonstrating accurate *α* recovery across the full prior range of demographic histories.

**Figure 14:**
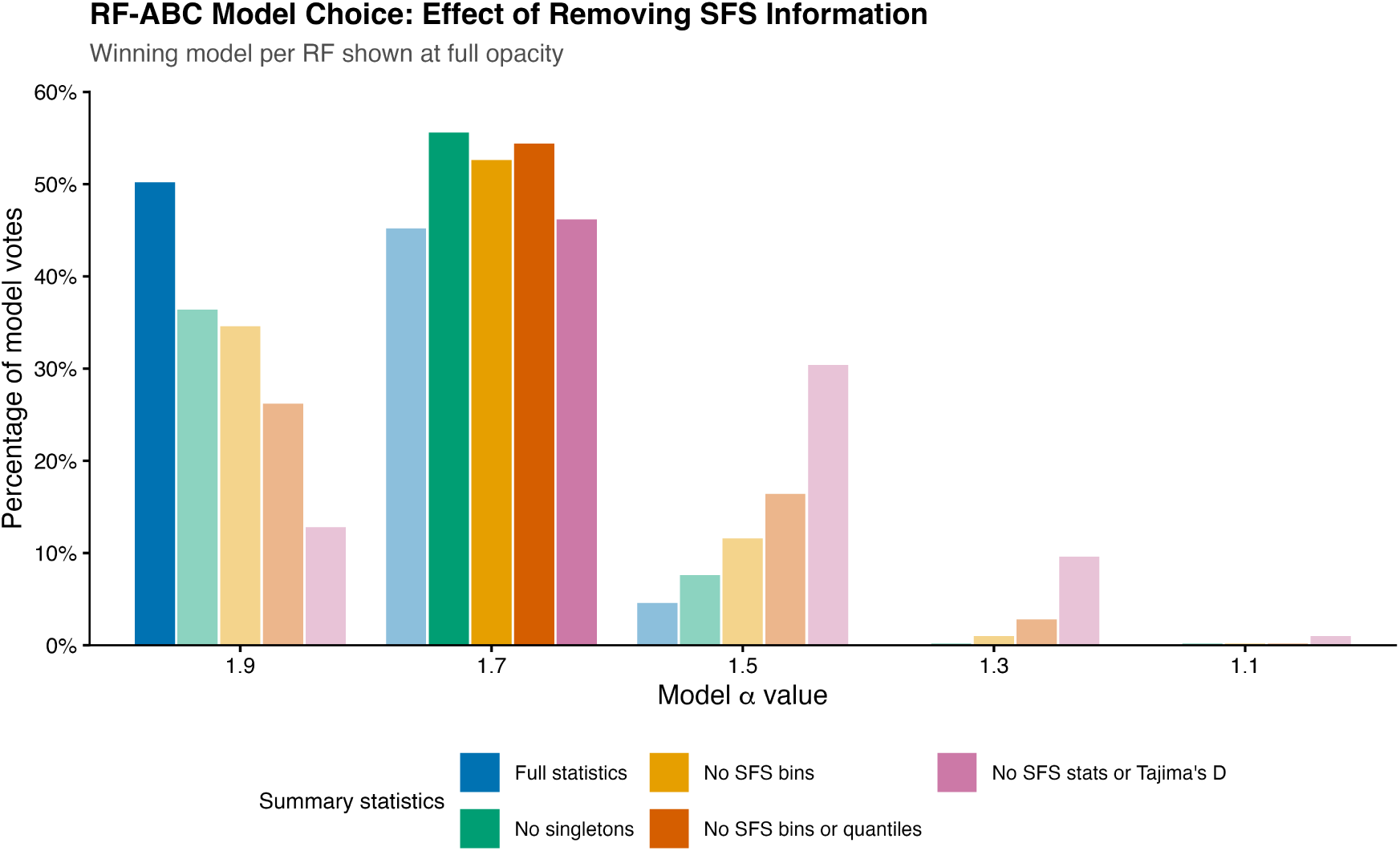
RF-ABC model choice for the *β*-coalescent parameter *α* across five progressively reduced summary statistic sets. Each bar shows the percentage of votes assigned to each *α* model (1.9, 1.7, 1.5, 1.3, 1.1) by a random forest trained on the indicated statistic subset; the winning model per random forest is shown at full opacity. When all statistics are included, the plurality of votes support *α* = 1.9, the conservative approximation of the Kingman coalescent, driven primarily by singleton frequency (fSFS1) and other SFS-derived statistics. As SFS information is progressively removed — first singletons, then all SFS bins, then AFS quantiles, and finally Tajima’s *D* — the plurality shifts to *α* = 1.7, a moderate multiple merger process. This pattern demonstrates that the initial Kingman-like signal is an artefact of demographic confounding in SFS-based statistics rather than a genuine signature of binary coalescence.

**Figure 15:**
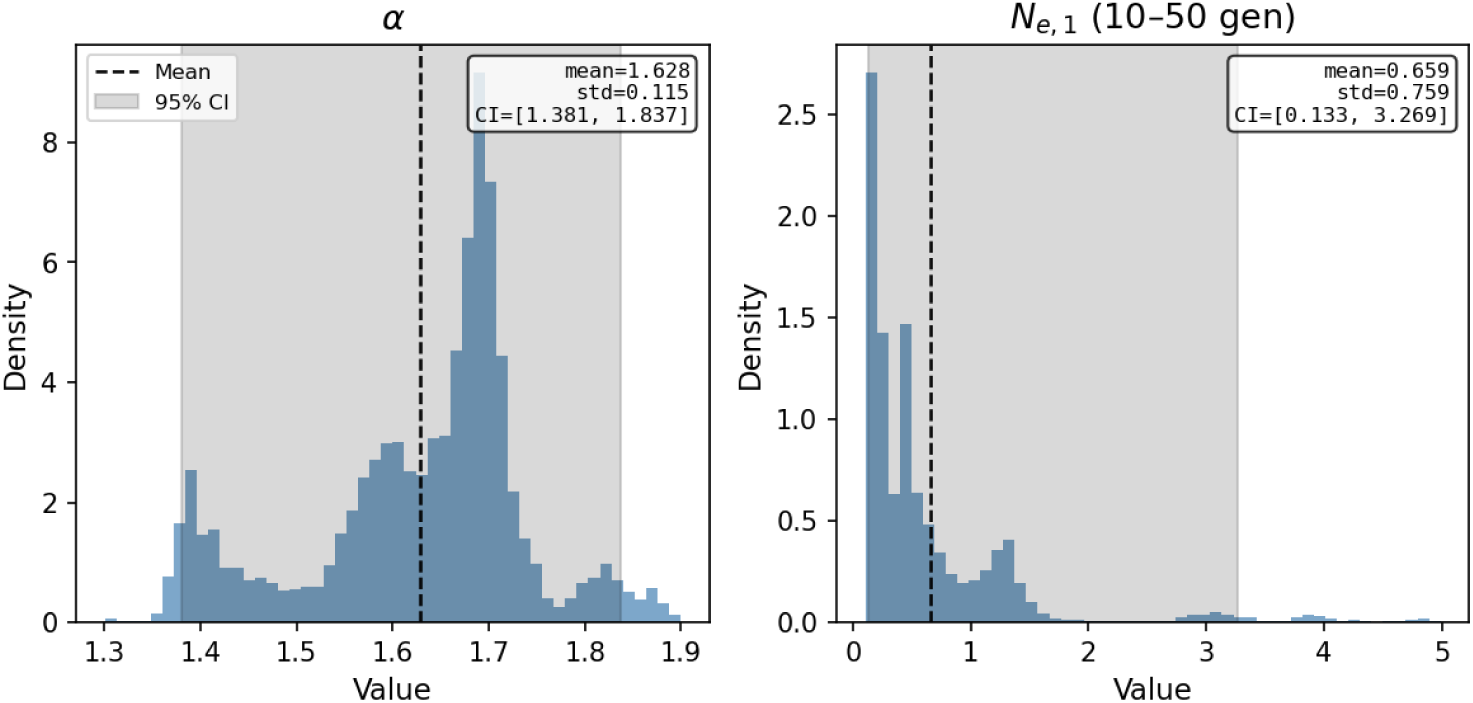
NSF posterior marginal distributions for the *β*-coalescent parameter *α* (left) and the most recent demographic window *N_e,_*_1_ (10–50 generations ago, right), based on 10,000 posterior samples conditioned on the observed summary statistics. Dashed line indicates the posterior mean; shaded band indicates the 95% credible interval. *α* is well-constrained with mean 1.628 ± 0.115 (95% CI [1.381, 1.837]), clearly excluding the Kingman limit (*α* = 1.9). *N_e,_*_1_ shows a multimodal distribution concentrated near low values (mean 0.659, median 0.441) but with a wide credible interval ([0.133, 3.269]), consistent with a recent population size reduction that is credible but not resolved with high confidence. The contrast between the tighter *α* posterior and the diffuse *N_e,_*_1_ posterior illustrates that the *β*-coalescent parameter is identifiable from the data while individual demographic windows are not.

**Figure 16:**
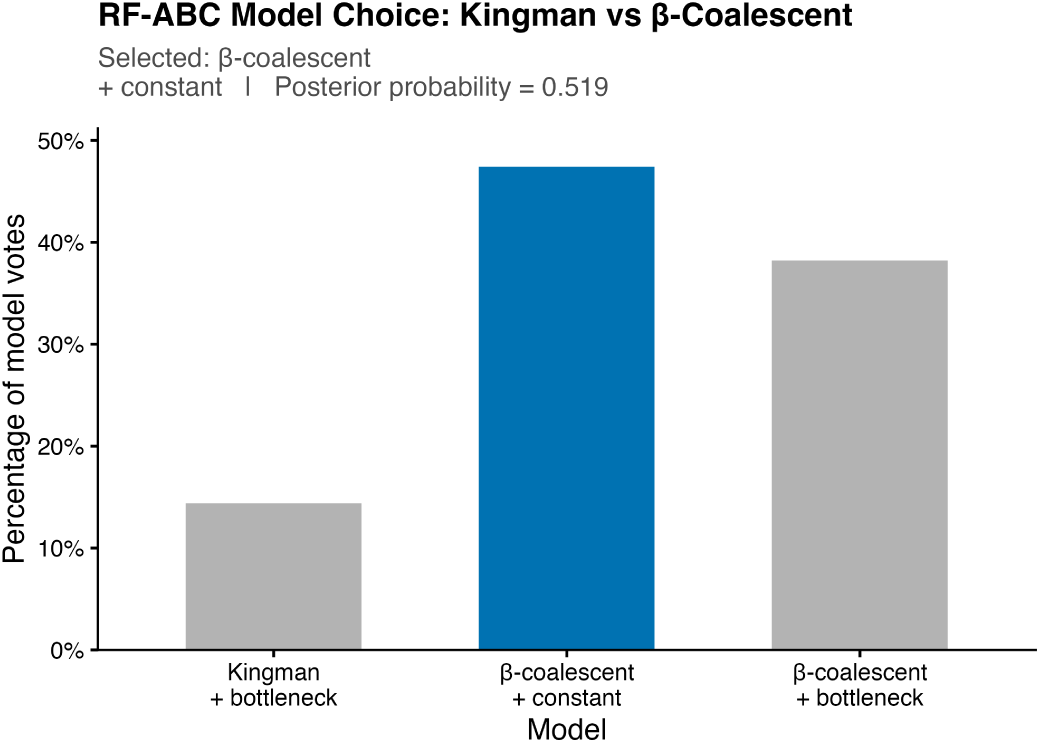
RF-ABC model choice between Kingman coalescence with a demographic bottleneck, *β*-coalescent with constant demography, and *β*-coalescent with a demographic bottleneck. The winning model is shown in blue. The *β*-coalescent with constant demography receives the plurality of votes (posterior probability = *post.prob*), providing evidence against a Kingman coalescent process.

## 4 Discussion

In this study, we first attempt to jointly assess multiple-merger coalescent (MMC) processes and changes in past population size in *Blumeria hordei*, thereby significantly advancing our understanding of coalescent dynamics in organisms with highly skewed reproductive success. Second, we attempt to dissect the underpinning causal mechanisms generating MMC signatures, namely between selective and neutral sweepstakes reproduction. Our findings have important implications for both population genomic methodology and our understanding of pathogen evolution.

### 4.1 Evidence for MMC in *B. hordei*

The Neural Posterior Estimation (NPE) framework confidently inferred a moderate multiple-merger process in *B. hordei* (*α* = 1.627 ± 0.116), distinct from the conservative Kingman approximation (*α* = 1.9). This finding aligns with our biological understanding of *B. hordei*’s life history characteristics, which may include highly skewed offspring distributions during sporulation, epidemic growth patterns, and recurrent bottlenecks associated with seasonal cycles and host availability. These life-history and reproductive features create ideal conditions for multiple-merger events, in which a small number of individuals contribute disproportionately to the next generation Waples et al. 2013; Tellier and Lemaire 2014; Waples et al. 2018; Eldon 2020; Árnason et al. 2023; Menardo, Gagneux, and Freund 2021.

The evidence for MMC in *B. hordei* parallels recent findings in the closely related *Blumeria graminis* f.sp. *tritici* (Jigisha et al. 2025), suggesting that multiple-merger processes may be a common feature of *Blumeria* species evolution. This taxonomic consistency strengthens the case for MMC as a fundamental aspect of powdery mildew population dynamics, with potential implications for understanding the evolutionary characteristics of other biotrophic plant pathogens with similar life history traits.

### 4.2 Demographic History of Crop Pathogens

The inferred demographic history, while requiring cautious interpretation due to wide confidence intervals, suggests a complex population trajectory that may reflect agricultural intensification. The best-fitting simulation showed population expansion around 300 years ago, which would coincide with the expansion of barley agriculture during the industrial revolution and pre-modern era, followed by a decline coinciding with the Green Revolution and widespread adoption of fungicides and resistance breeding. While we cannot definitively attribute these patterns to anthropogenic factors without additional evidence, the timing is consistent with major changes in agricultural practices that could have dramatically altered *B. hordei* population dynamics.

This potential demographic response to agricultural intensification highlights the dynamic interactions among pathogen populations, recent human activities, and disease management. Understanding these historical demographic patterns is crucial for predicting future evolutionary trajectories and developing sustainable disease management strategies. The integration of MMC processes with demographic modeling provides a powerful framework for interpreting these patterns and their implications for pathogen evolution, whereas previous studies have relied on inferring demography or MMC, without accounting for the confounding effects between these processes and their signatures.

### 4.3 Underpinnings of MMC and Selection Detection

The presence of MMC processes in *B. hordei* has significant implications for detecting genuine selective signatures. Multiple-merger genealogies can generate patterns that mimic or mask signals of positive selection (Schweinsberg 2003; Durrett and Schweinsberg 2004; Tellier and Lemaire 2014), potentially leading to false positives or negatives when standard selection detection methods are applied (Korfmann et al. 2024a). Our demonstration that demographic history can substantially alter SFS characteristics under MMC (with singleton frequencies differing by up to 10% between constant and variable population sizes) underscores the importance of accounting for both coalescent model specification and demographic complexity in selection scans. We have shown the robustness of RAiSD and SweeD to neutral MMC and demographic history by adjusting the significance threshold, as recommended in state-of-the-art population genomics analyses (Johri et al. 2022). However, this yields a counter-intuitive result. Using a conservative approach (taking the top 0.1% outlier regions) we found few signatures of sweeps (1,600 with RaisD and 120 with SweeD) and a high threshold for RaisD of 246.48. However, when the threshold was informed conservatively by the simulations under MMC and demography, had a value of 8 and yielded 189,597 putative sweep regions (that is 11.6 % of all considered genomic regions). We interpret this results as an indication that MMC in is due to two processes. First, a neutral mechanism likely due to biological and life-history traits (clonal/sexual phases and Boom-and-Bust cycles), which could explain a genome-wide background level of MMC signature (*α* value of ca. 1.6). Second, when correcting for this background “noise” of MMC, we find still a large number of regions under potential positive selective pressure, suggesting the occurrence of pervasive genome-wide selection. Such selection could occur at genes involved in infectivity (for example, effector genes) to overcome barley resistance, or at genes favoring resistance to fungicides. Nonetheless, we cannot exclude that the genetic draft caused by pervasive positive selection (Achaz and Schertzer 2023) could generate part of the neutral MMC signatures we detect. Another possibility is the occurrence of genome-wide pervasive background selection, which can bias genealogies across the genome (Johri, Charlesworth, and Jensen 2020) and, under low to zero recombination, may appear as widespread positive selection signatures (Stephan 2010). The latter may be especially seen in centromeric regions.

The development of MMC-aware selection validation represents a crucial next step for understanding adaptive evolution in *B. hordei* and similar organisms. Approaches like SweeD and RAisD, which rely on the SFS, likely require recalibration that explicitly accounts for multiple-merger processes with demography. This is particularly important for agriculturally relevant pathogens where accurate identification of adaptive loci is essential for understanding virulence evolution and resistance breakdown (Jigisha et al. 2025).

### 4.4 Methodological Advances in Coalescent Inference

Our results highlight critical limitations of current population genomic methods when applied to organisms with complex evolutionary histories. Site frequency spectrum (SFS)-based statistics taken in isolation proved highly sensitive to demographic fluctuations, leading to systematic biases in coalescent inference. The random forest ABC analysis initially supported a Kingman-like process (*α* = 1.9) when all SFS information was included, but progressively shifted toward moderate MMC (*α* = 1.7) as SFS-based statistics were removed. This pattern demonstrates that apparent Kingman-like signals can be artifacts of demographic confounding rather than genuine signatures of binary coalescence.

The demographic inference results from MSMC2 further illustrate this methodological challenge. When applied to the observed *B. hordei* data, MSMC2 inferred an extreme demographic history with over 99% of lineages entering a recent bottleneck followed by rapid expansion. This artificial demographic pattern is a known property of Kingman model misspecification. In addition, this demography is discordant with the observed SFS characteristics and demonstrates how misspecified coalescent models can generate misleading demographic reconstructions.

Our NPE framework successfully addresses these limitations by inferring coalescent in the presence of complex demographic parameters while explicitly accounting for recombination effects. The method’s robustness is demonstrated by its high correlation (*r* = 0.81) between true and inferred *α* values across diverse demographic scenarios in simulation-based calibration tests. Importantly, while the NPE showed modest sensitivity to intermediate-timescale population size changes (50– 300 generations ago), this confounding effect explained less than 4% of the variance in inferred *α*, indicating that the method successfully disentangles MMC signals from the modeled demographic effects.

### 4.5 Implications for Population Genomics of Pathogens

Our findings extend beyond plant pathology to broader questions in population genomics of species with sweepstakes reproduction. The methodological advances presented here, particularly the NPE framework and the demonstration of SFS confounding, have applications for any organism characterized by highly variable reproductive success, demographic complexity, and recombination. Marine organisms with high fecundity (Waples et al. 2013; Waples et al. 2018; Árnason et al. 2023; Goldberg 2026), microbial populations (Freund et al. 2023; Menardo, Gagneux, and Freund 2021), and species experiencing rapid range expansions may all benefit from MMC-aware analytical approaches.

Several limitations warrant discussion. First, our sample is geographically restricted to Germany, limiting conclusions about global *B. hordei* population structure and MMC prevalence across the species range. Expanding sampling to include diverse geographical populations would strengthen inferences about the generality of MMC processes in this species. Second, while our NPE framework successfully accounts for demographic complexity, the wide confidence intervals for historical population sizes indicate limited power to resolve fine-scale demographic details from current summary statistics.

Future research should focus on developing selection detection methods that explicitly account for MMC processes, expanding geographical sampling to assess global patterns of coalescent variation, and integrating additional data types (such as structural variants and transposable elements) that may provide complementary insights into *B. hordei* evolution. The framework developed here also provides a foundation for comparative studies of MMC across related species, potentially revealing fundamental patterns in the evolution of biotrophic plant pathogens.

## Data availability

All raw sequence data are deposited in the publicly accessible European Nucleotide Archive (ENA) under accession number PRJEB112051. The scripts for analysing and plotting the results have been published on GitLab of the Prof. for Population Genetics (XXXXX)

## Funding

This project was funded by the German Research Foundation (DFG) TRR 356/1 2023 – Project 491090170, subproject A03 to Ralph Hückelhoven and Auŕelien Tellier.

## Acknowledgment

We would like to thank Veronika Schmitt and Africa Fabro Espada from the Chair of Phytopathology, TUM, and Daniela Scheikl from the Division of Population Genetics, TUM, in handling of *B. hordei* isolates. We also thank Daniela Scheikl for expert DNA purification. Furthermore, we thank EpiLogic GmbH Agrarbiol. Forschung und Beratung (Freising, Germany) and colleagues for the provision of *B. hordei* isolates. We further acknowledge project I01 of the TRR PlantMicrobe for assistance with data management, including the implementation of FAIR data principles and the facilitation of data and code sharing.

## 6 Supplementary Tables

**Supplementary Table 1:** List of the 54 German *Bh* isolates used in this study. Headers: Group refers to the sample collection by year and federal state; LAT and LON refer to the approximate latitude and longitude of the sample; PLZ is the German post code recorded; Date is the sampling date; Ro Route: if given, sampling tour during which the sample was taken, and LAT and LON were then evenly assigned along the route to the samples. In all other cases, LAT and LON report the actual sample location. Abbreviations: *^c^* clonal with another line; *^s^*^2^ or *^s^*^3^ sub-population 2 or 3; *^L^* ONT long-read sequencing was conducted; ^∗^ low output for *^L^*. R1 = route Freising to Plattling, R2 = route Plattling to Regensburg to Waidhaus, R3 = route Rothenburg to Schweinfurt to Hammelburg. M = MAT: mating type, either 1 = MAT-1-1-1 or 2 = MAT-1-2-1

| Isolate | Group | LAT | LON | PLZ | Date | Ro | M |
| --- | --- | --- | --- | --- | --- | --- | --- |
| TUM1<sup>L</sup> | Ref | 48.3998 | 11.7216 | 85354 | - |  | 1 |
| Bay43<sup>L</sup> | 23_Bav | 48.9788 | 11.0333 | 91788 | 2023-05-19 |  | 1 |
| Bay50<sup>L</sup> | 23_Bav | 48.4006 | 11.6837 | 85354 | 2023-06-09 |  | 2 |
| Bay52<sup>c</sup> | 23_Bav | 48.4006 | 11.6837 | 85354 | 2023-06-09 |  | 2 |
| Bay53 | 24_Bav | 48.0027 | 12.6403 | 83349 | 2024-04-29 |  | 2 |
| Bay54 | 24_Bav | 48.4687 | 11.9257 | 85368 | 2024-06-03 |  | 1 |
| Bay55 | 24_Bav | 47.8452 | 12.0917 | 83026 | 2024-05-08 |  | 1 |
| DE01_01 | 24_Bav | 48.3892 | 11.7471 | 85354 | 2024-05-14 | R1 | 1 |
| DE01_02 | 24_Bav | 48.4151 | 11.9586 | 85354 | 2024-05-14 | R1 | 1 |
| DE01_03<sup>s3,L*</sup> | 24_Bav | 48.4218 | 11.9001 | 85354 | 2024-05-14 | R1 | 2 |
| DE01_04<sup>s3,L*</sup> | 24_Bav | 48.4781 | 12.1125 | 85354 | 2024-05-14 | R1 | 1 |
| DE01_05 | 24_Bav | 48.5261 | 12.1871 | 85354 | 2024-05-14 | R1 | 1 |
| DE01_06 | 24_Bav | 48.6147 | 12.2771 | 85354 | 2024-05-14 | R1 | 1 |
| DE01_07 | 24_Bav | 48.6090 | 12.4023 | 85354 | 2024-05-14 | R1 | 2 |
| DE01_08 | 24_Bav | 48.6150 | 12.4825 | 85354 | 2024-05-14 | R1 | 1 |
| DE01_09<sup>c</sup> | 24_Bav | 48.6763 | 12.6732 | 85354 | 2024-05-14 | R1 | 1 |
| DE01_10<sup>s3</sup> | 24_Bav | 48.6661 | 12.7724 | 85354 | 2024-05-14 | R1 | 2 |
| DE01_11<sup>c</sup> | 24_Bav | 48.8306 | 12.8441 | 85354 | 2024-05-14 | R1 | 2 |
| DE01_12 | 24_Bav | 48.7453 | 12.8476 | 94447 | 2024-05-14 | R2 | 2 |
| DE02_01 | 24_Bav | 48.7481 | 12.7088 | 94447 | 2024-05-14 | R2 | 1 |
| DE02_02 | 24_Bav | 48.8782 | 12.5869 | 94447 | 2024-05-14 | R2 | 1 |
| DE02_03<sup>c</sup> | 24_Bav | 48.8233 | 12.3471 | 94447 | 2024-05-14 | R2 | 1 |
| DE02_04 | 24_Bav | 49.0044 | 12.2197 | 94447 | 2024-05-14 | R2 | 2 |
| DE02_05 | 24_Bav | 49.0868 | 12.0903 | 94447 | 2024-05-14 | R2 | 1 |
| DE02_06 | 24_Bav | 48.9899 | 12.0247 | 94447 | 2024-05-14 | R2 | 2 |
| DE02_07 | 24_Bav | 49.1071 | 12.1371 | 94447 | 2024-05-14 | R2 | 1 |
| DE02_08 | 24_Bav | 49.2377 | 12.2147 | 94447 | 2024-05-14 | R2 | 2 |
| DE02_09 | 24_Bav | 49.3708 | 12.2776 | 94447 | 2024-05-14 | R2 | 2 |
| DE02_10 | 24_Bav | 49.4908 | 12.5004 | 94447 | 2024-05-14 | R2 | 2 |
| DE02_11 | 24_Bav | 49.7058 | 12.5224 | 94447 | 2024-05-14 | R2 | 2 |
| DE02_12<sup>L</sup> | 24_Bav | 49.6416 | 12.4944 | 94447 | 2024-05-14 | R2 | 2 |
| DE03_01<sup>L</sup> | 24_Bav | 49.4477 | 10.2043 | 91541 | 2024-05-28 | R3 | 2 |
| EL01_01<sup>L</sup> | 23_Bra | 53.1135 | 11.8443 | 19348 | 2023-03-22 |  | 1 |
| EL01_02<sup>s2</sup> | 23_Bra | 53.1135 | 11.8443 | 19348 | 2023-03-23 |  | 1 |
| EL02_02<sup>L</sup> | 23_Bra | 52.2112 | 14.6374 | 15295 | 2023-03-23 |  | 2 |
| EL03_01 | 23_Bra | 52.1938 | 13.2842 | 14959 | 2023-03-24 |  | 2 |
| EL03_02<sup>s2</sup> | 23_Bra | 52.1938 | 13.2842 | 14959 | 2023-03-24 |  | 1 |
| EL04_01 | 23_Bra | 52.3481 | 14.5063 | 15234 | 2023-05-08 |  | 2 |
| EL04_02 | 23_Bra | 52.3481 | 14.5063 | 15234 | 2023-05-08 |  | 1 |
| FS01<sup>L</sup> | 23_Bav | 48.4056 | 11.6927 | 85354 | 2023-05-02 |  | 1 |
| FS02<sup>c</sup> | 23_Bav | 48.4113 | 11.7398 | 85354 | 2023-06-16 |  | 1 |
| FS03 | 23_Bav | 48.4113 | 11.7398 | 85354 | 2023-06-19 |  | 1 |
| FS04 | 23_Bav | 48.3914 | 11.7375 | 85354 | 2023-07-14 |  | 2 |
| FS05 | 23_Bav | 48.3914 | 11.7375 | 85354 | 2023-07-17 |  | 1 |
| FS07 | 23_Bav | 48.3908 | 11.7693 | 85356 | 2023-07-22 |  | 1 |
| FS08 | 24_Bav | 48.3998 | 11.7216 | 85354 | 2024-04-16 |  | 2 |
| FS09 | 24_Bav | 48.4051 | 11.7525 | 85354 | 2024-04-29 |  | 2 |
| FS10 | 24_Bav | 48.3998 | 11.7216 | 85354 | 2024-04-16 |  | 2 |
| FS11 | 24_Bav | 48.4018 | 11.7243 | 85354 | 2024-05-25 |  | 1 |
| FS12 | 24_Bav | 48.3908 | 11.7693 | 85356 | 2024-06-03 |  | 1 |
| FS13<sup>L</sup> | 24_Bav | 48.3914 | 11.7375 | 85354 | 2024-06-12 |  | 1 |
| FS14 | 24_Bav | 48.3914 | 11.7375 | 85354 | 2024-06-12 |  | 1 |
| Muc01<sup>L</sup> | 23_Bav | 48.1413 | 11.5602 | 80355 | 2023-07-17 |  | 1 |
| Muc02<sup>L</sup> | 24_Bav | 48.1413 | 11.5602 | 80356 | 2024-06-06 |  | 2 |

**Supplementary Table 2:**
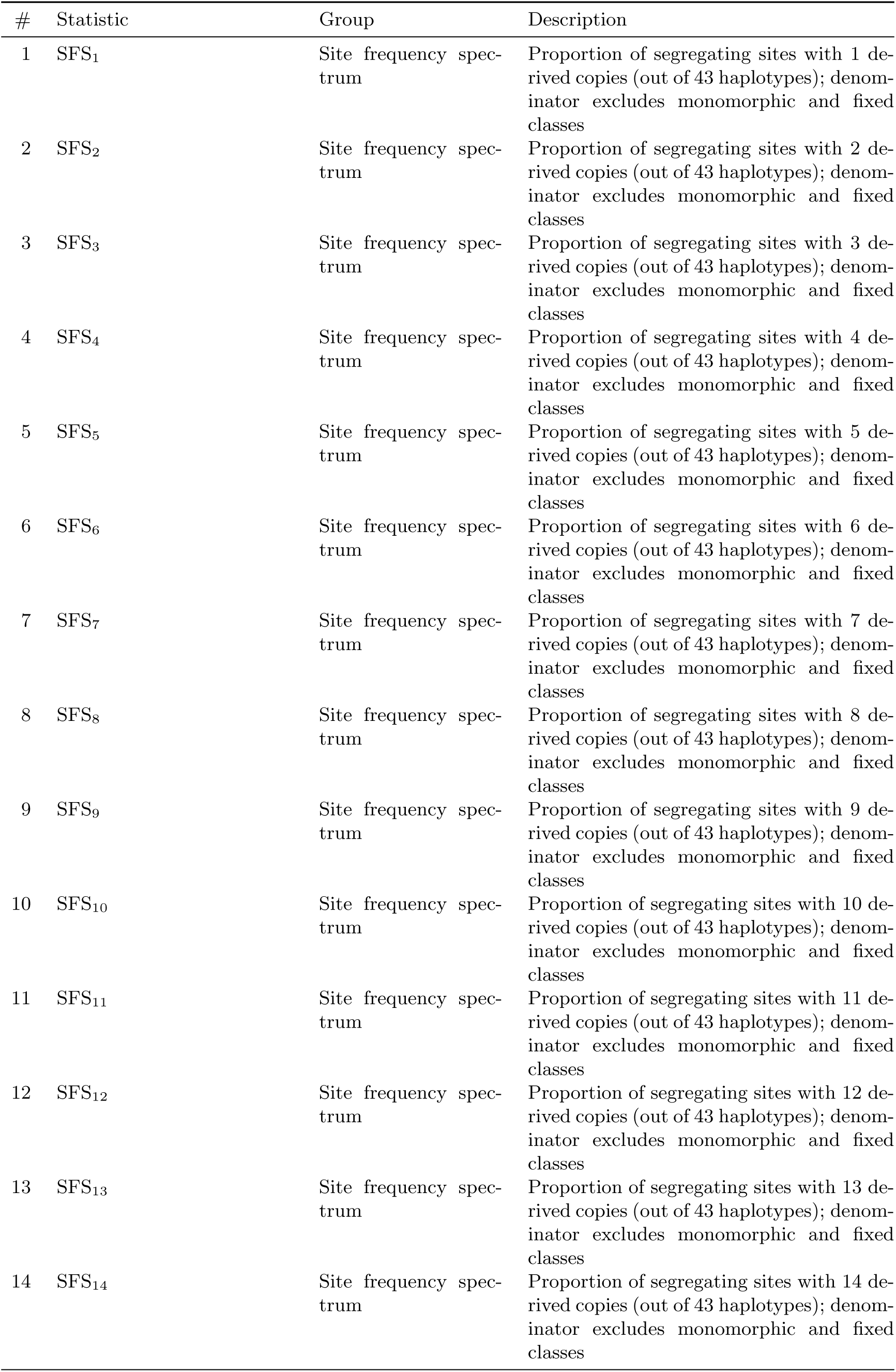
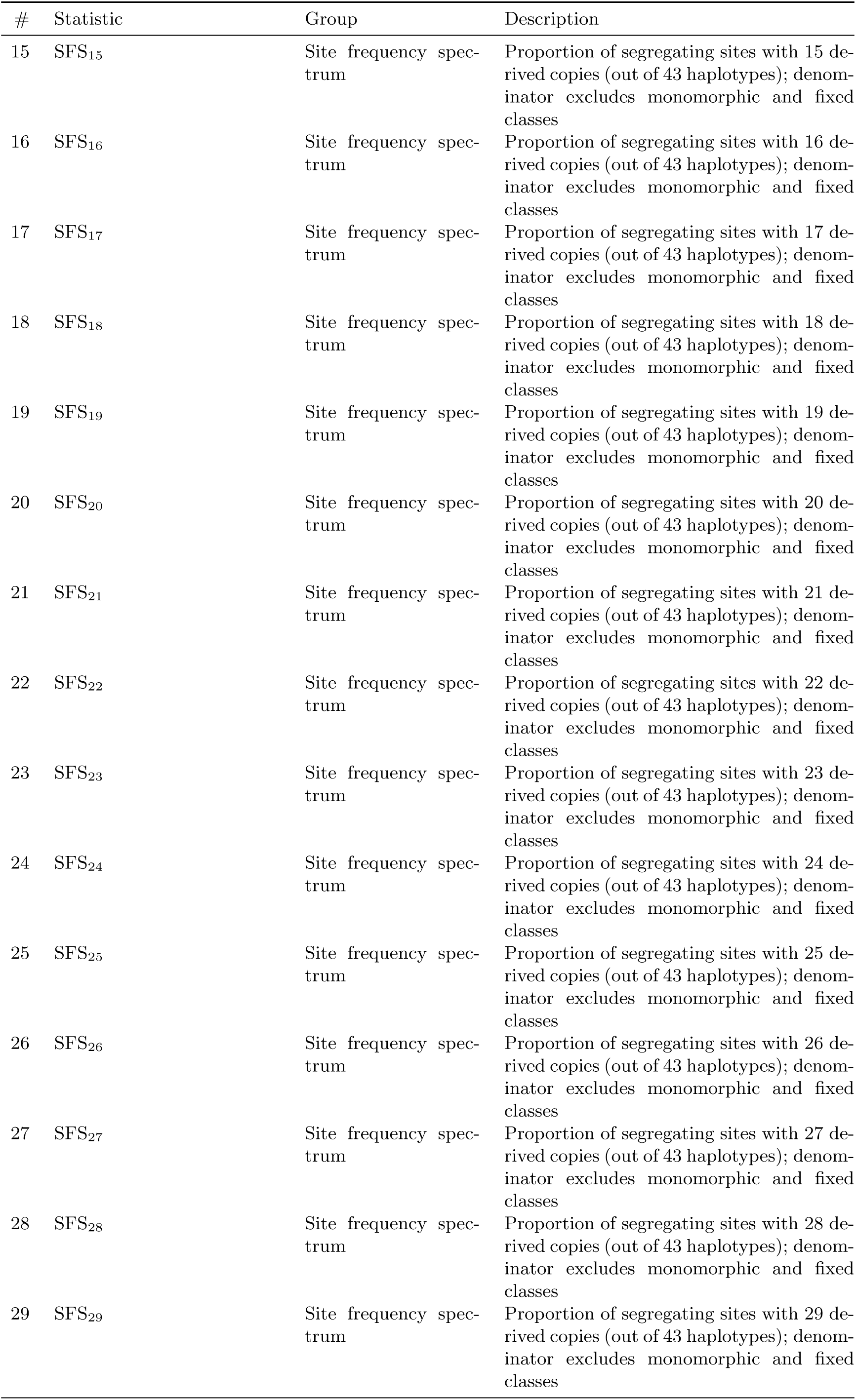
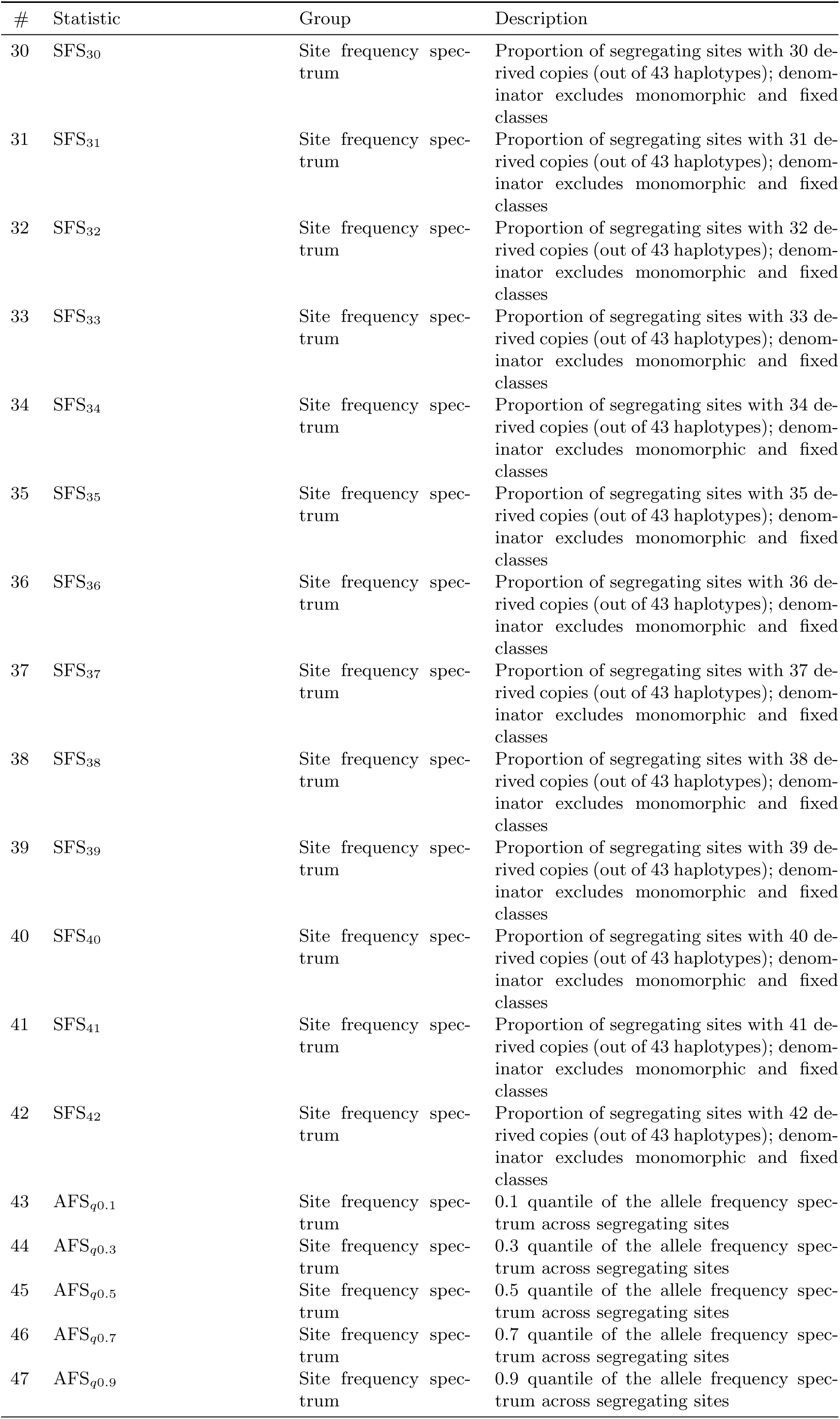
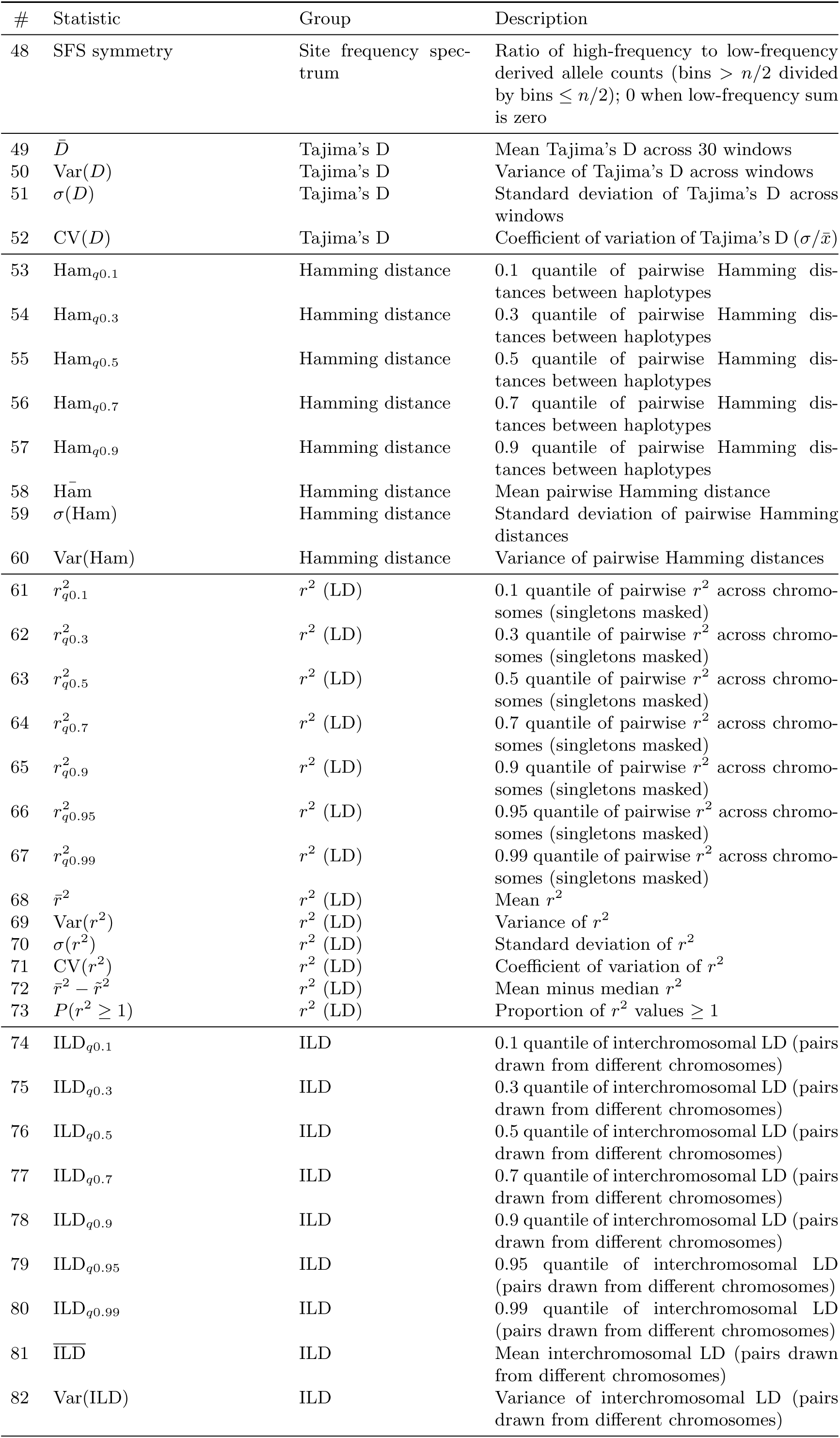
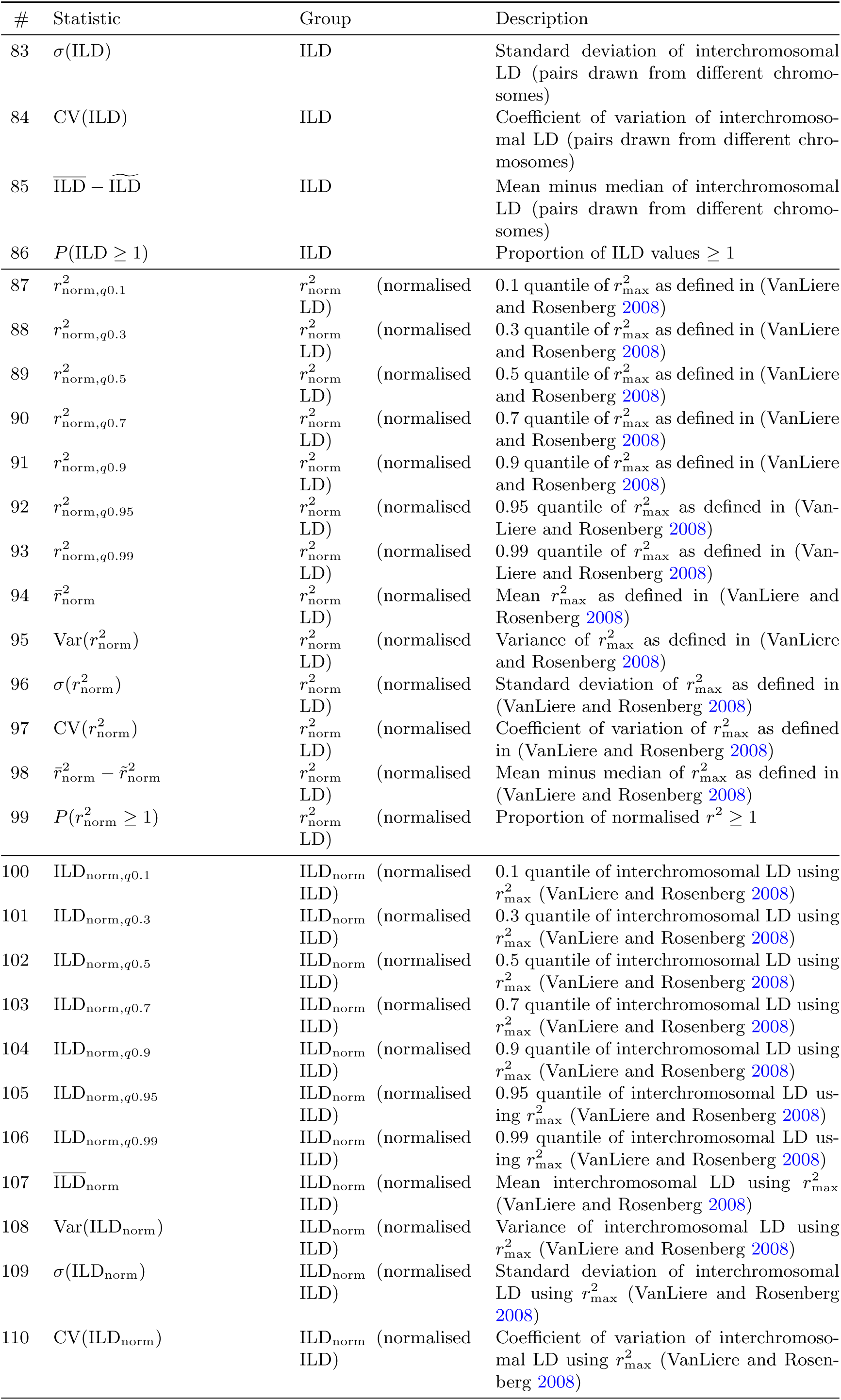
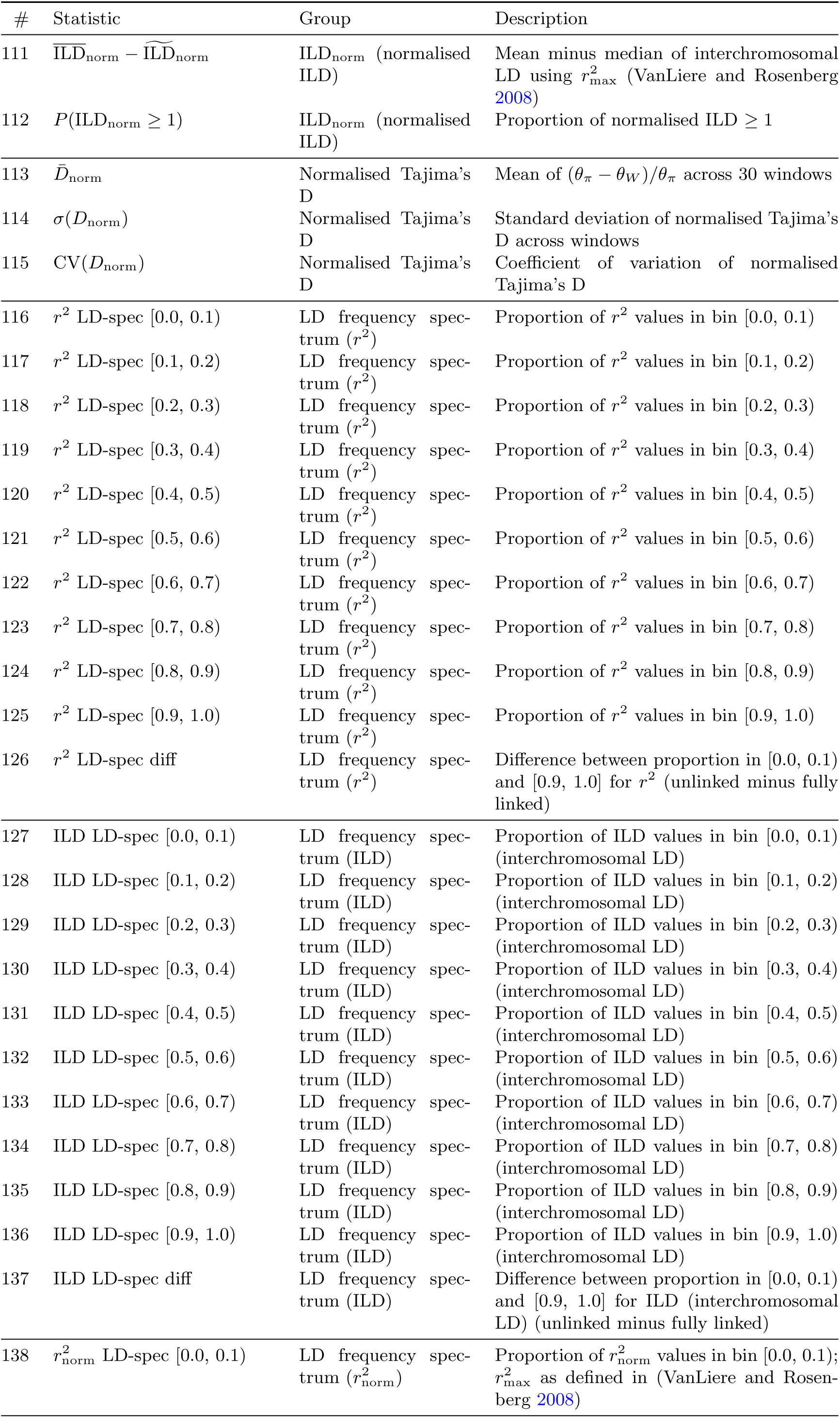
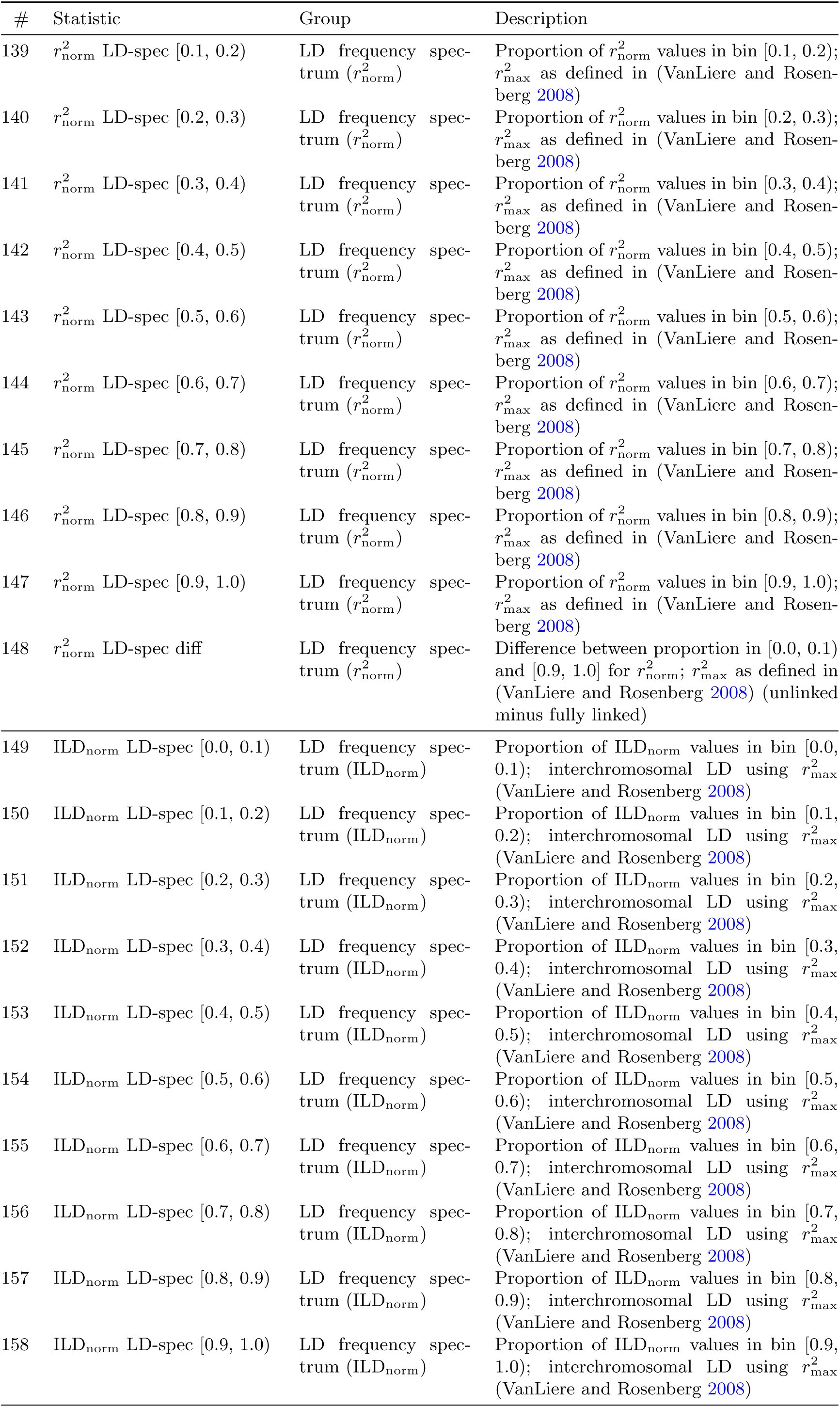
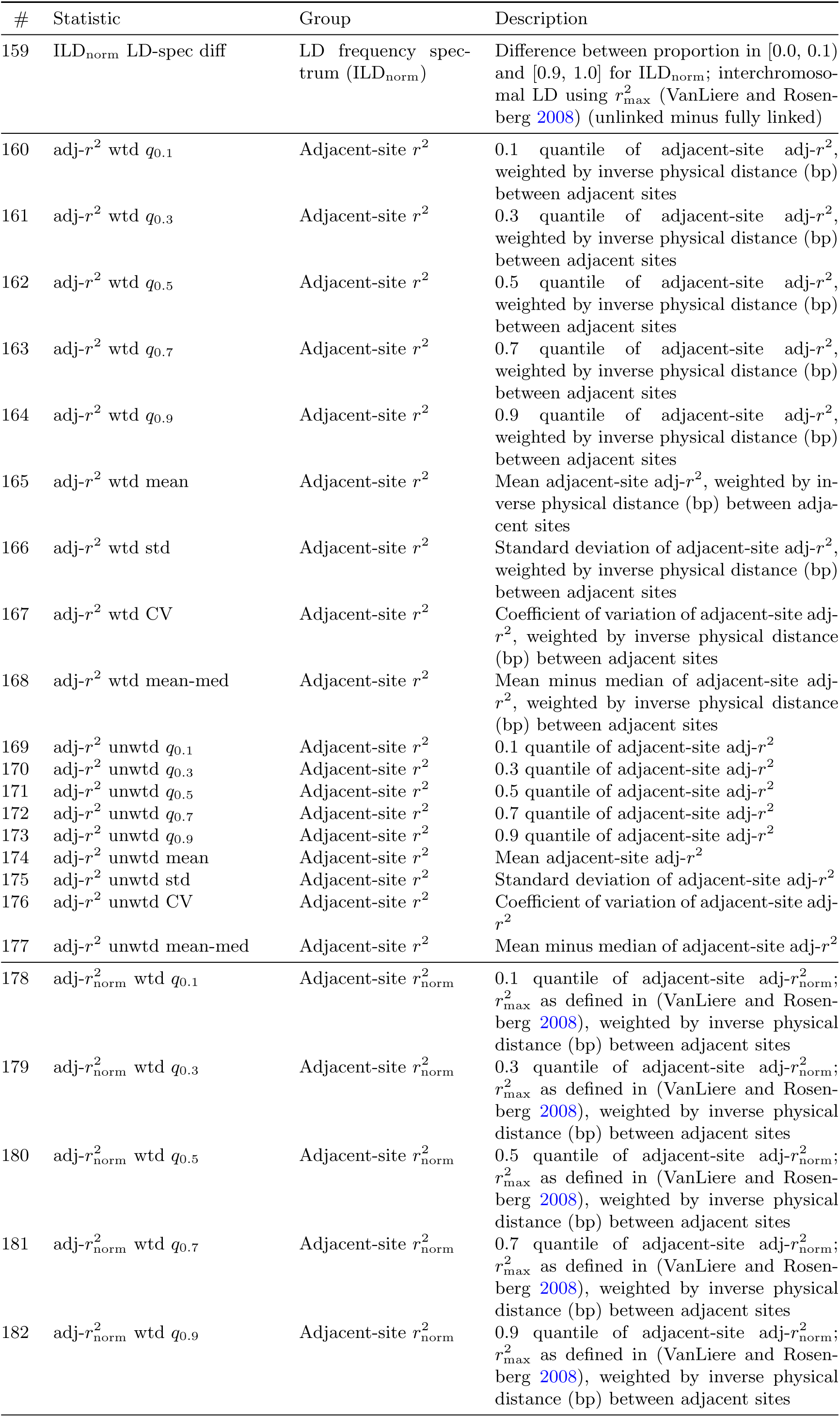
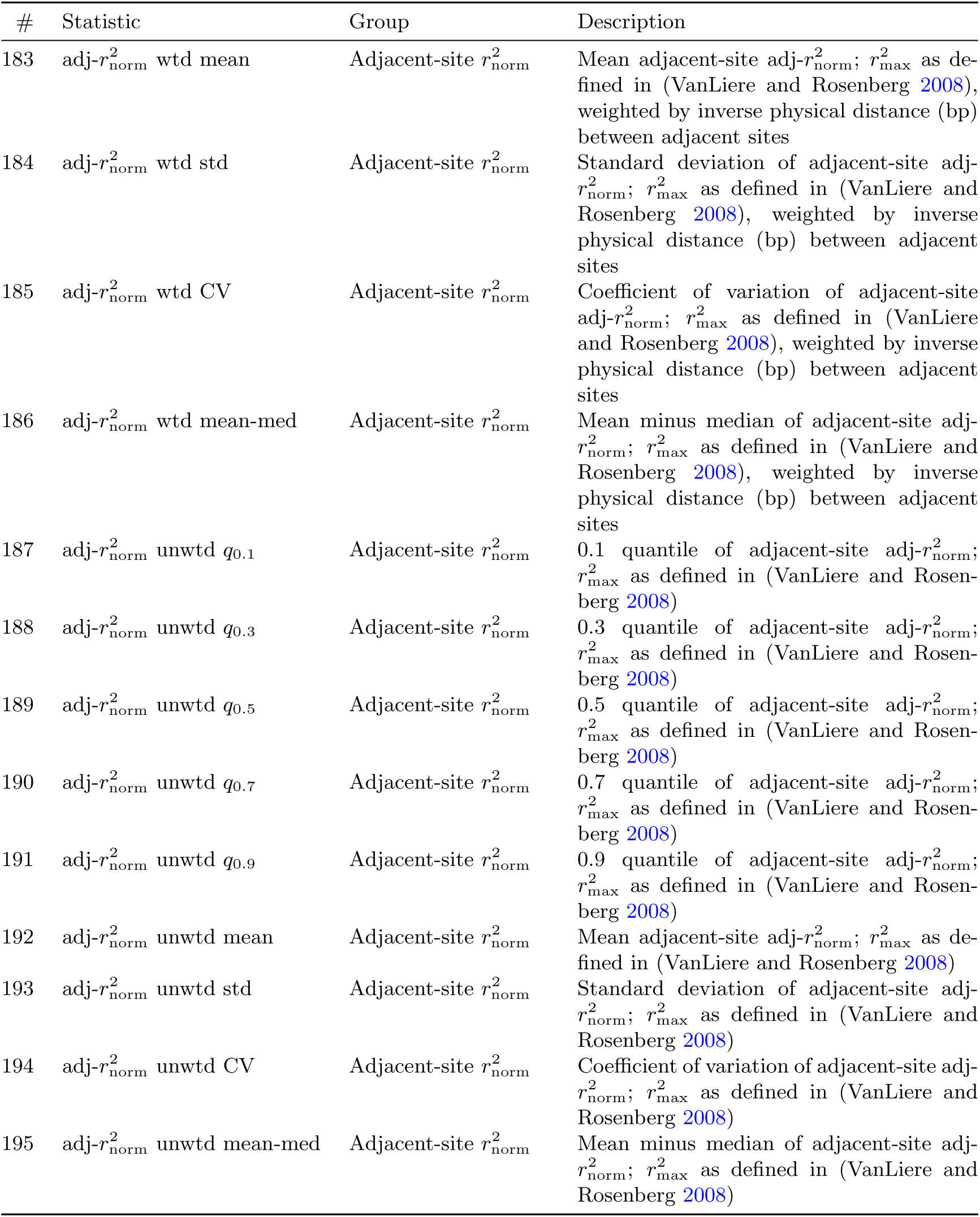
Summary statistics used in simulation-based inference. All 195 statistics are computed for each simulated and observed dataset. *n* = 43 haplotypes; singletons are masked for LD statistics. CV = *σ*/x̅.

**Supplementary Figure 1:**
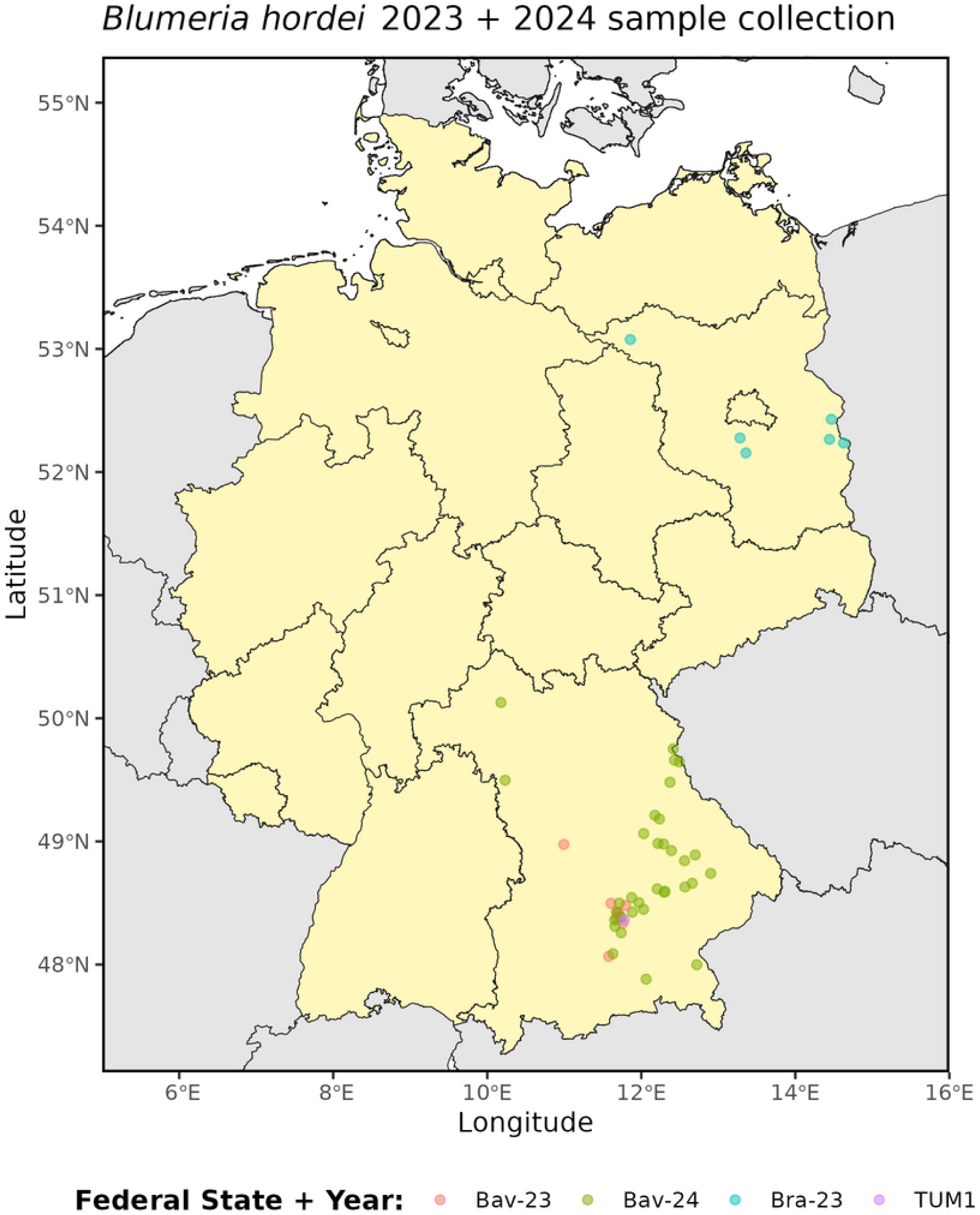
Map of Germany with approximate collection points of 53 *B. hordei* isolates and the reference. Colour of points indicate the federal state and year of sampling. TUM1 is the reference isolate.

**Supplementary Figure 2:**
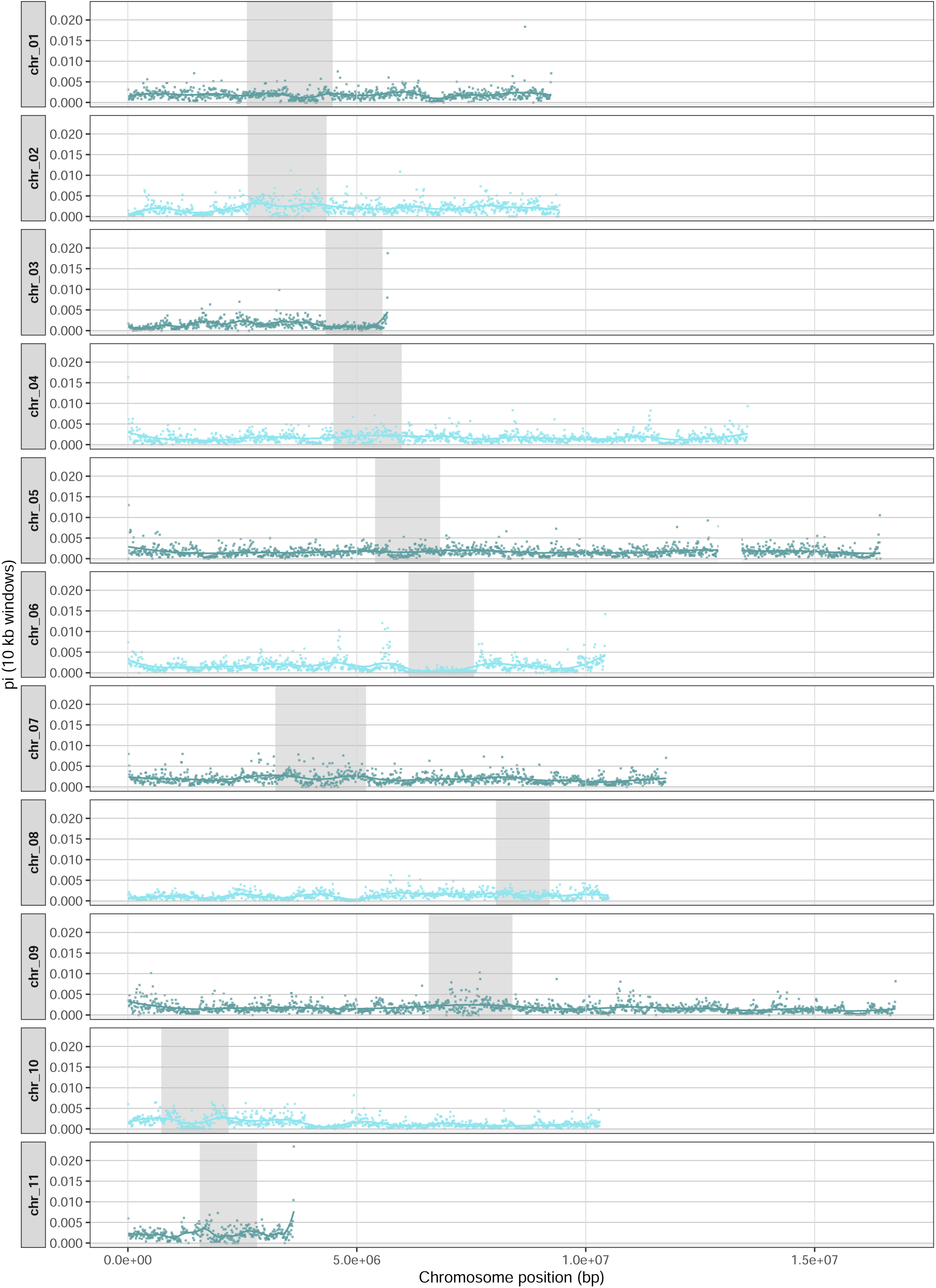
Nucleotide diversity *pi* along the 11 chromosomes of *Bh*; calculated in 10 Mb windows. The centromer region is indicated by a dark grey shading. The two parts of chr 05 are shown in one panel, with chr 05a on the left and chr 05b on the right from the white gap. Units are needed.

**Supplementary Figure 3:**
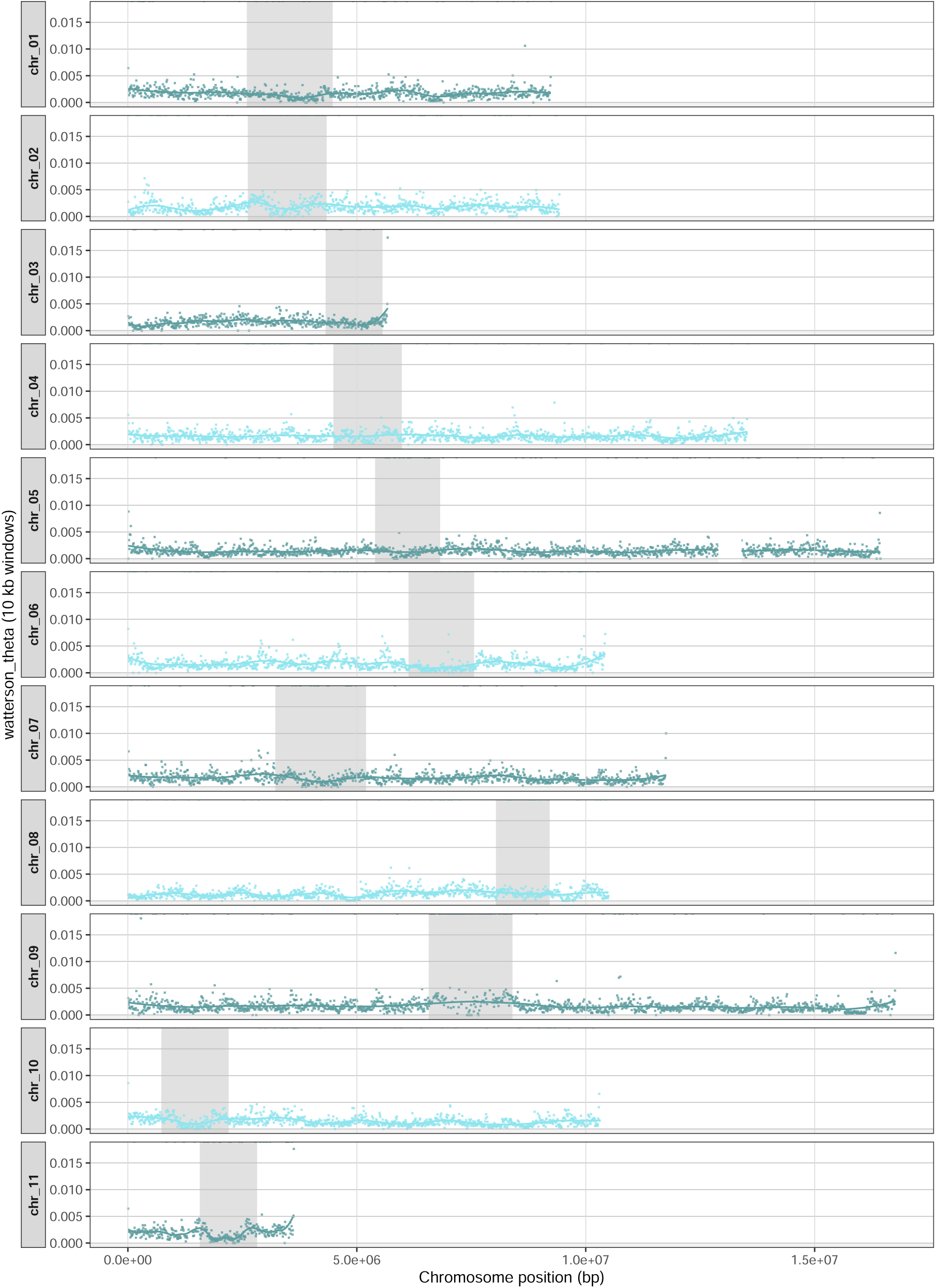
Watterson’s *theta* along the 11 chromosomes of *Bh*; calculated in 10 Mb windows. The centromer region is indicated by a dark grey shading. The two parts of chr 05 are shown in one panel, with chr 05a on the left and chr 05b on the right from the white gap. Units are needed.

**Supplementary Figure 4:**
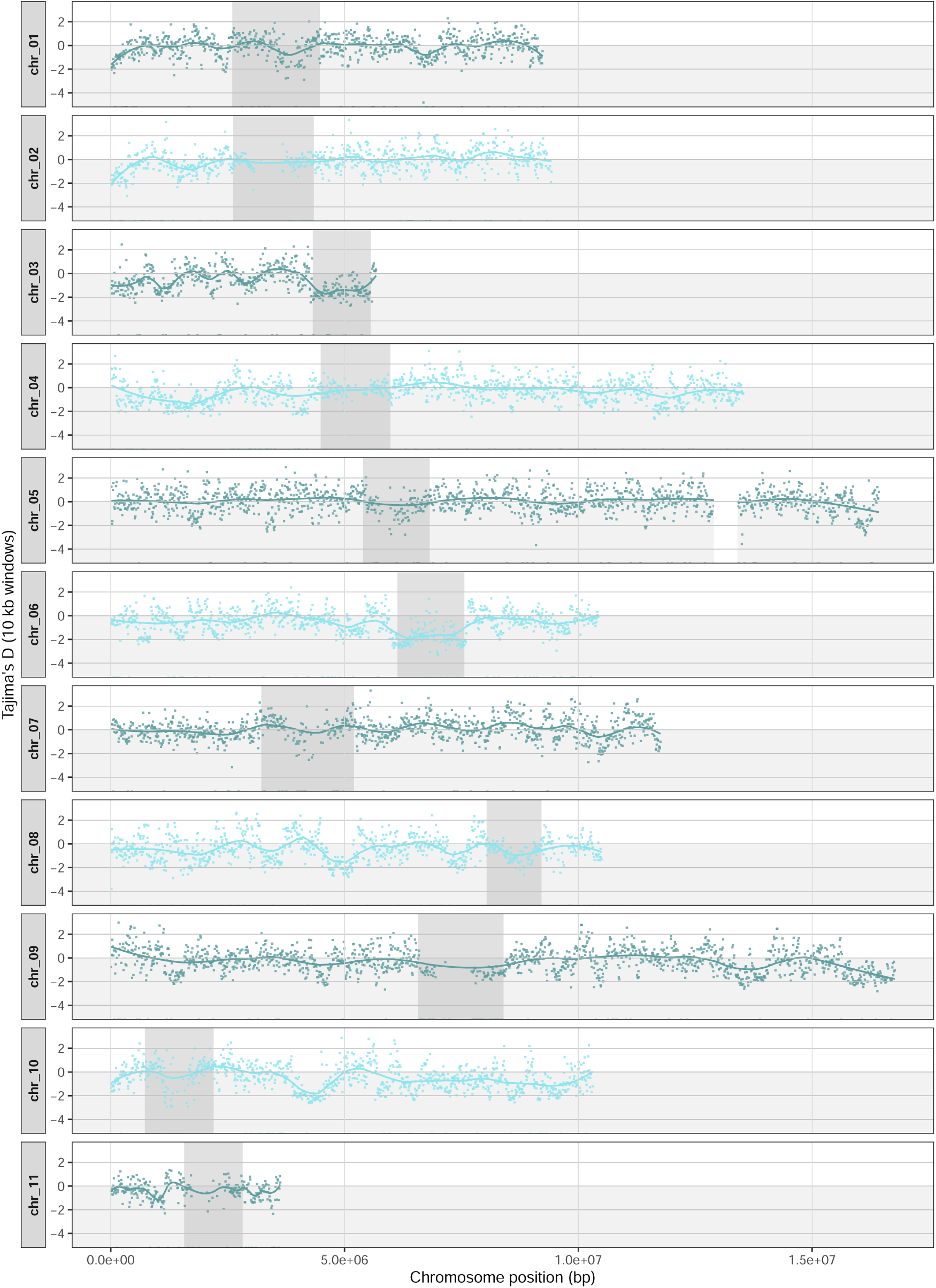
Tajima’s D along the 11 chromosomes of *Bh*; calculated in 10 Mb windows. The centromer region is indicated by a dark grey shading. The two parts of chr 05 are shown in one panel, with chr 05a on the left and chr 05b on the right from the white gap. Units are needed.

**Supplementary Figure 5:**
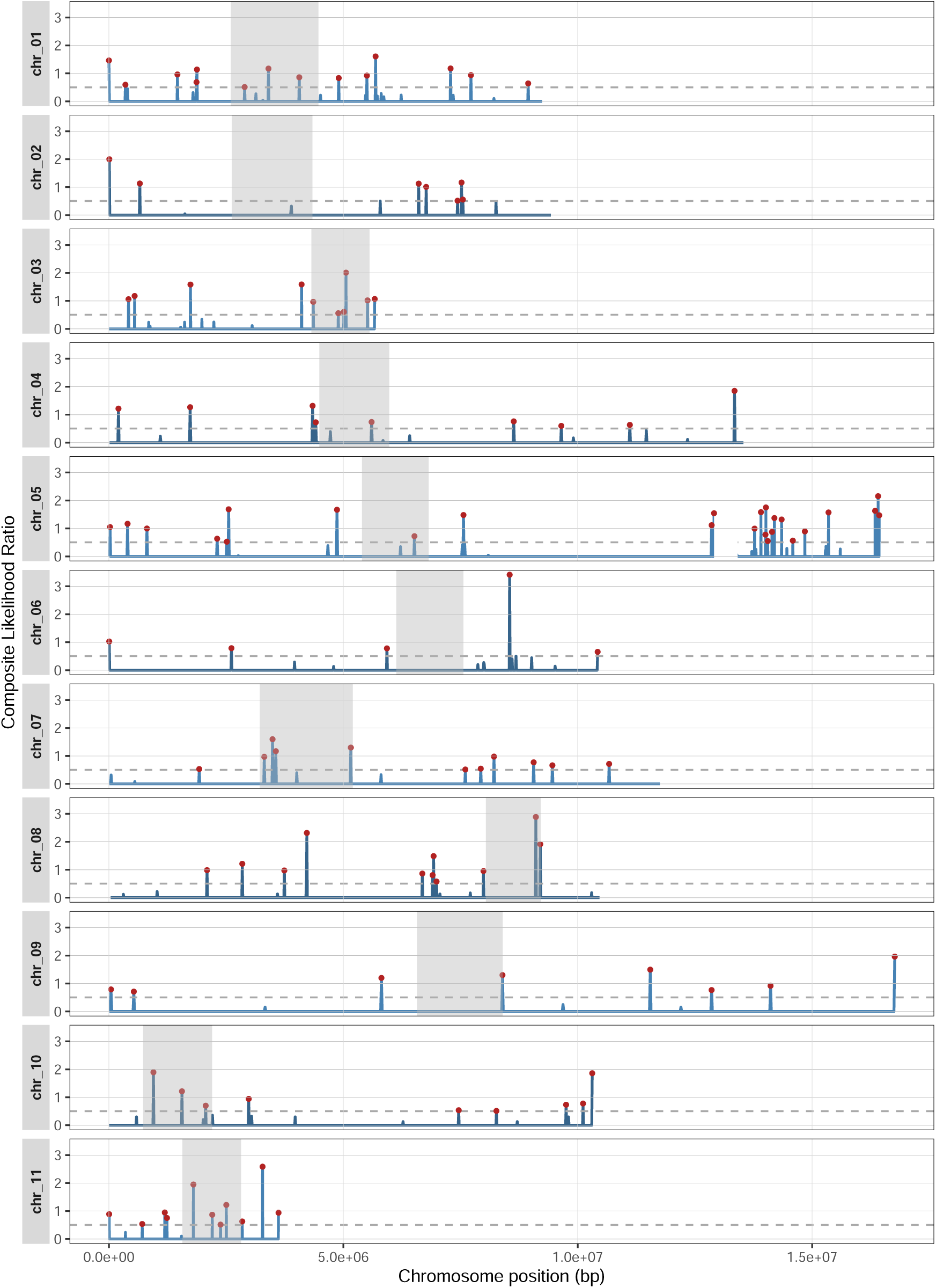
Scans for positive selection along the 11 chromosomes of *Bh* using SweeDv4.0 conducted on the short-read data of 43 isolates. The Composite Likelihood Ratio (CLR) is given in blue, with the 1% top sweeps marked in red. The dashed horizontal lines indicate the 99% significance threshold (CLR = 0.50). The centromere regions are indicated by a dark grey shading. The two parts of chr 05 are shown in one panel, with chr 05a on the left and chr 05b on the right of the white gap.

**Supplementary Figure 6:**
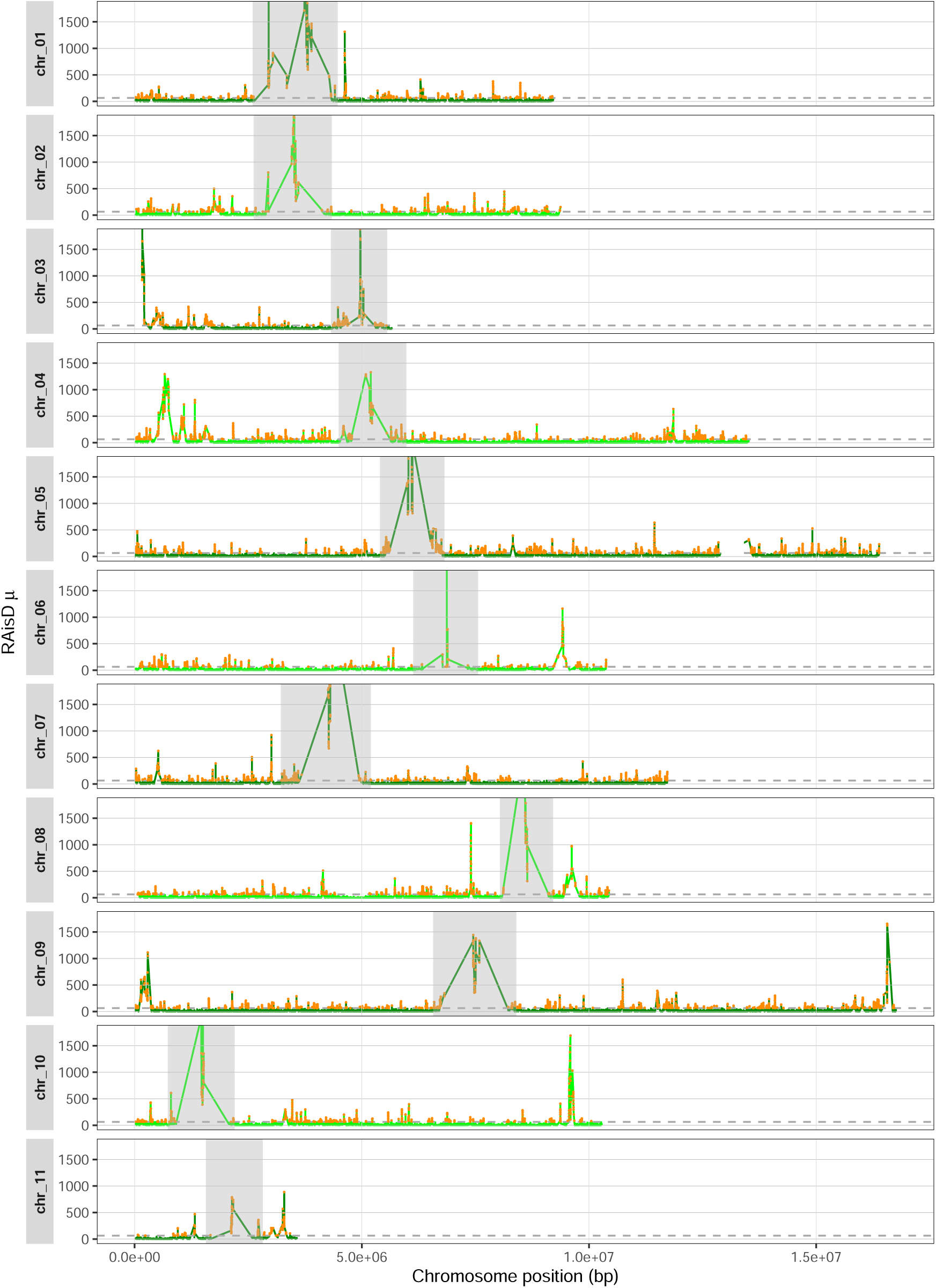
Scans for positive selection along the 11 chromosomes of *Bh* using RAiSD conducted on the short-read data of 43 isolates. The RAiSD *µ* score is given in green, with the 0.1% top sweeps marked in orange (1631 regions). The dashed horizontal lines indicate the 99.9% significance threshold (*µ_thr_*= 246.48). The y-axis is capped at *µ* = 1800 to show details of smaller sweeps. The full plots are available in Fig S7. The centromere regions are indicated by a dark grey shading. The two parts of chr 05 are shown in one panel, with chr 05a on the left and chr 05b on the right of the white gap.

**Supplementary Figure 7:**
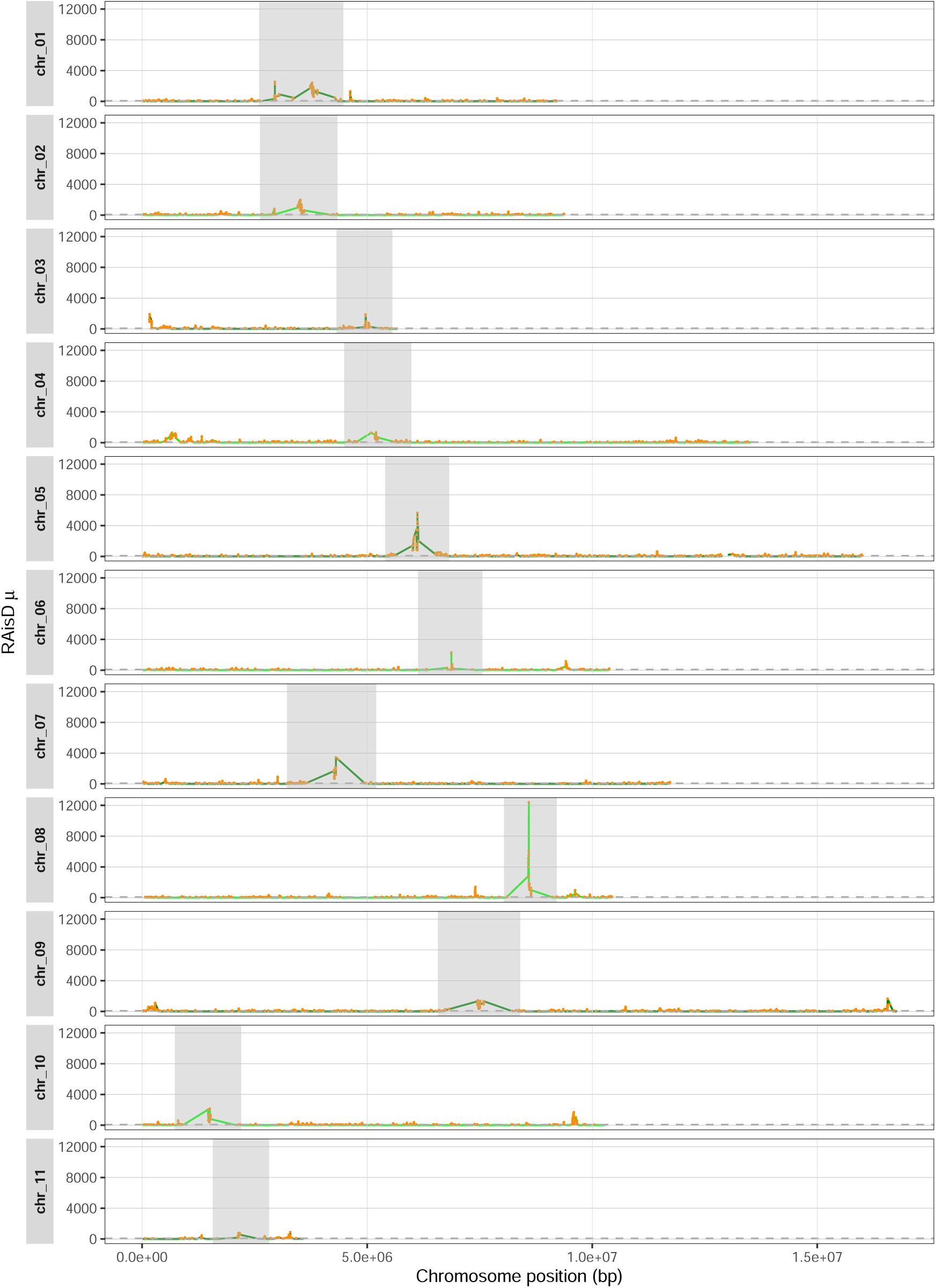
Scans for positive selection along the 11 chromosomes of *Bh* using RAiSD conducted on the short-read data of 43 isolates. The RAiSD *µ* score is given in green, with the 0.1% top sweeps marked in orang (1631 regions). The dashed horizontal lines indicate the 99.9% significance threshold (*µ_thr_* = 246.48). The centromere regions are indicated by a dark grey shading. The two parts of chr 05 are shown in one panel, with chr 05a on the left and chr 05b on the right of the white gap.

**Supplementary Figure 8:**
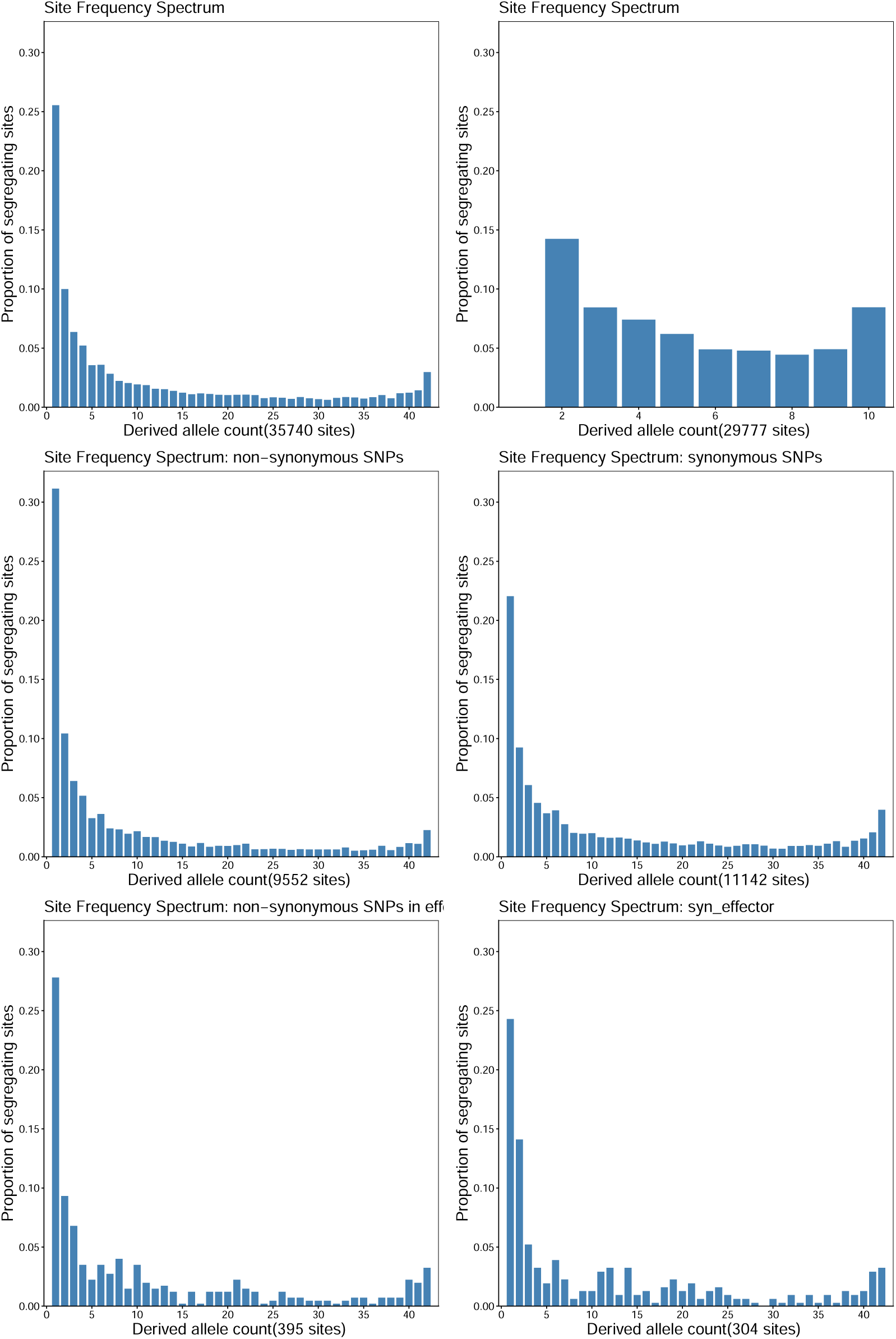
Folded SFS of in the *Bh* population. All segregating sites using short-read data from 43 isolates (top left) or long-read data from 11 isolates (top right). Using the short-read data, the SFS was also calculated for non-synonymous and synonymous sites: Non-synonymous (middle left) and synonymous (middle right) sites of all genes. Non-synonymous (middle left) and synonymous (middle right) sites of the 874 hypothetical effector genes.

**Supplementary Figure 9:**
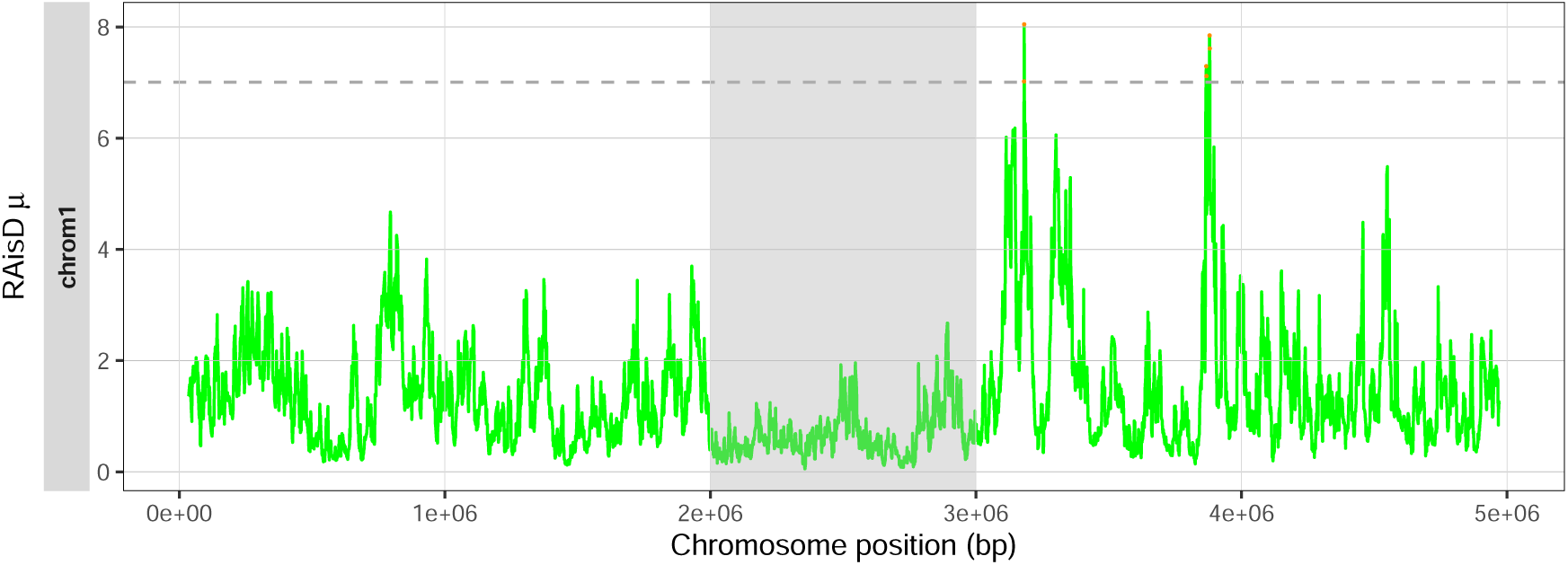
Validation of RAiSD for selective sweep detection on neutral *β*-coalescent with demography. *µ* values are consistently below 8, suggesting robusteness against signals caused purely by neutral and moderate multiple merger coalescence.

**Supplementary Figure 10:**
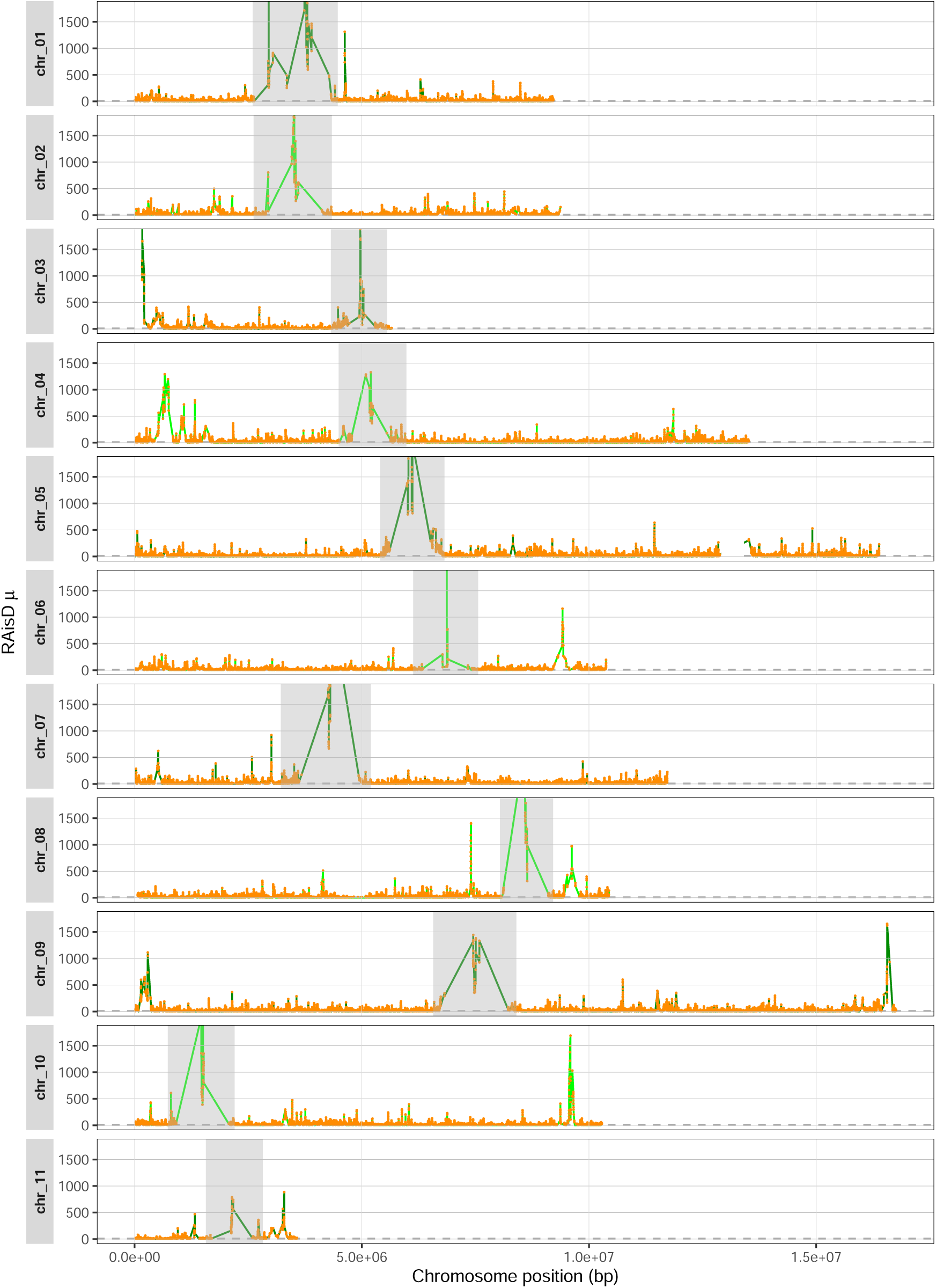
Scans for positive selection along the 11 chromosomes of *Bh* using RAiSD conducted on the short-read data of 43 isolates. A total of 189,597 sweep regions (11.82 % of all regions) are identified, if the RAiSD *µ* threshold of 8.0, taken from the scan of the simulated data using netural *β*-coalescent with demography (Figure S9), is applied. The *µ* score is given in green, with sweeps above the threshold marked in orange. The centromere regions are indicated by a dark grey shading. The two parts of chr 05 are shown in one panel, with chr 05a on the left and chr 05b on the right of the white gap.

## Notes

### Competing Interest Statement

The authors have declared no competing interest.

## References

Achaz, G and E Schertzer (2023). “Weak genetic draft and the Lewontin’s paradox”. In: bioRxiv.

Agrios, George N (2024). Agrios’ Plant Pathology. Academic Press.

Alachiotis, Nikolaos and Pavlos Pavlidis (June 2018). “RAiSD detects positive selection based on multiple signatures of a selective sweep and SNP vectors”. In: Communications Biology 1.1, p. 79. issn: 2399-3642. doi: 10.1038/s42003-018-0085-8. url: https://doi.org/10.1038/s42003-018-0085-8.

Arendt, Elke K. and Emanuele Zannini (2013). “4 - Barley”. In: Cereal Grains for the Food and Beverage Industries. Ed. by Elke K. Arendt and Emanuele Zannini. Woodhead Publishing Series in Food Science, Technology and Nutrition. Woodhead Publishing, 155–201e. isbn: 978-0-85709-413-1. doi: 10.1533/9780857098924.155. url: https://www.sciencedirect.com/science/article/pii/B9780857094131500048.

Árnason, Einar et al. (2023). “Sweepstakes reproductive success via pervasive and recurrent selective sweeps”. In: Elife 12, e80781.

Bailey, Nick, Laurie Stevison, and Kieran Samuk (2025). “Correcting for Bias in Estimates of *θ_w_* and Tajima’s D From Missing Data in Next-Generation Sequencing”. In: Molecular Ecology Resources 25.6. e14104 MER-24-0389.R1, e14104. doi: 10.1111/1755-0998.14104. eprint: https://onlinelibrary.wiley.com/doi/pdf/10.1111/1755-0998.14104. url: https://onlinelibrary.wiley.com/doi/abs/10.1111/1755-0998.14104.

Baumdicker, Franz, et al. (Dec. 2021). “Efficient ancestry and mutation simulation with msprime 1.0”. In: Genetics 220.3, iyab229. issn: 1943-2631. doi: 10.1093/genetics/iyab229. eprint: https://academic.oup.com/genetics/article-pdf/220/3/iyab229/43780247/iyab229.pdf. url: https://doi.org/10.1093/genetics/iyab229.

Bindschedler, Laurence V., Ralph Panstruga, and Pietro D. Spanu (Feb. 2016). “Mildew-Omics: How Global Analyses Aid the Understanding of Life and Evolution of Powdery Mildews”. In: Frontiers in Plant Science 7. Publisher: Frontiers Media SA. issn: 1664-462X. doi: 10.3389/fpls.2016.00123. url: http://journal.frontiersin.org/Article/10.3389/fpls.2016.00123/abstract.

Birkner, Matthias, Jochen Blath, and Bjarki Eldon (2013). “An ancestral recombination graph for diploid populations with skewed offspring distribution”. In: Genetics 193.1, pp. 255–290.

Brown, J. K. M. and A. C. Jessop (1995). “Genetics of avirulences in Erysiphe graminis f.sp. hordei”. In: Plant Pathology 44.6, pp. 1039–1049. doi: 10.1111/j.1365-3059.1995.tb02663.x. eprint: https://bsppjournals.onlinelibrary.wiley.com/doi/pdf/10.1111/j.1365-3059.1995.tb02663.x. url: https://bsppjournals.onlinelibrary.wiley.com/doi/abs/10.1111/j.1365-3059.1995.tb02663.x.

Brown, James KM et al. (2002). “Oases in the desert: dispersal and host specialization of biotrophic fungal pathogens of plants”. In: Dispersal ecology, pp. 395–409.

Bushnell, B. (2014). BBTools GitHub repository. https://github.combbushnell/BBTools. Official website: https://bbmap.org.

Chang, Christopher C et al. (Dec. 2015). “Second-generation PLINK: rising to the challenge of larger and richer datasets”. In: GigaScience 4.1, s13742–015–0047–8. issn: 2047-217X. doi: 10.1186/s13742-015-0047-8. eprint: https://academic.oup.com/gigascience/article-pdf/4/1/s13742-015-0047-8/60713591/gigascience_4_1_s13742-015-0047-8.pdf. url: https://doi.org/10.1186/s13742-015-0047-8.

Cingolani, Pablo et al. (2012). “A program for annotating and predicting the effects of single nucleotide polymorphisms, SnpEff”. In: Fly 6.2, pp. 80–92. doi: 10.4161/fly.19695.

Danecek, Petr, et al. (Feb. 2021). “Twelve years of SAMtools and BCFtools”. In: GigaScience 10.2, giab008. issn: 2047-217X. doi: 10.1093/gigascience/giab008. eprint: https://academic.oup.com/gigascience/article-pdf/10/2/giab008/60687743/giab008.pdf. url: https://doi.org/10.1093/gigascience/giab008.

Dreiseitl, Antonın (2019). “Great pathotype diversity and reduced virulence complexity in a Central European population of Blumeria graminis f. sp. hordei in 2015–2017”. In: European Journal of Plant Pathology 153.3, pp. 801–811.

Dreiseitl, Antonın (2024). “Mlo-Mediated Broad-Spectrum and Durable Resistance against Powdery Mildews and Its Current and Future Applications”. In: Plants 13.1. issn: 2223-7747. doi: 10.3390/plants13010138. url: https://www.mdpi.com/2223-7747/13/1/138.

Durrett, Richard and Jason Schweinsberg (2004). “Approximating selective sweeps”. In: Theoretical population biology 66.2, pp. 129–138.

Eldon, Bjarki (2020). “Evolutionary genomics of high fecundity”. In: Annual Review of Genetics 54.1, pp. 213–236.

Eldon, Bjarki, et al. (Mar. 2015). “Can the Site-Frequency Spectrum Distinguish Exponential Population Growth from Multiple-Merger Coalescents?” In: Genetics 199.3, pp. 841–856. issn: 1943-2631. doi: 10.1534/genetics.114.173807. eprint: https://academic.oup.com/genetics/article-pdf/199/3/841/42133781/genetics0841.pdf. url: https://doi.org/10.1534/genetics.114.173807.

Ellwood, Simon R., Francisco J. Lopez-Ruiz, and Kar-Chun Tan (2024). “Barley powdery mildew control in Western Australia and beyond”. In: Plant Pathology 73.7, pp. 1666–1674. doi: 10.1111/ppa.13884. eprint: https://bsppjournals.onlinelibrary.wiley.com/doi/pdf/10.1111/ppa.13884. url: https://bsppjournals.onlinelibrary.wiley.com/doi/abs/10.1111/ppa.13884.

Frantzeskakis, Lamprinos, Stefan Kusch, and Ralph Panstruga (2018). “The need for speed: compartmentalized genome evolution in filamentous phytopathogens”. In: Molecular plant pathology 20.1, p. 3.

Frantzeskakis, Lamprinos, et al. (Dec. 2018). “Signatures of host specialization and a recent transposable element burst in the dynamic one-speed genome of the fungal barley powdery mildew pathogen”. In: BMC Genomics 19.1. Publisher: Springer Science and Business Media LLC. issn: 1471-2164. doi: 10.1186/s12864-018-4750-6. url: https://bmcgenomics.biomedcentral.com/articles/10.1186/s12864-018-4750-6.

Freund, Fabian and Arno Siri-Jegousse (2021). “The impact of genetic diversity statistics on model selection between coalescents”. In: Computational Statistics and Data Analysis 156, p. 107055. issn: 0167-9473. doi: 10.1016/j.csda.2020.107055. url: https://www.sciencedirect.com/science/article/pii/S0167947320301468.

Freund, Fabian, et al. (Mar. 2023). “Interpreting the pervasive observation of U-shaped Site Frequency Spectra”. In: PLOS Genetics 19, e1010677. doi: 10.1371/journal.pgen.1010677.

Goldberg, Amy (2026). “Rare variation in malaria parasites biases population-genetic inference”. In: bioRxiv, pp. 2026–01.

Jigisha, Jigisha et al. (May 2025). “Population genomics and molecular epidemiology of wheat powdery mildew in Europe”. In: PLOS Biology 23.5. Ed. by Sophien Kamoun, e3003097. issn: 1545-7885. doi: 10.1371/journal.pbio.3003097. url: https://doi.org/10.1371/journal.pbio.3003097.

Johri, Parul, Brian Charlesworth, and Jeffrey D Jensen (2020). “Toward an evolutionarily appropriate null model: jointly inferring demography and purifying selection”. In: Genetics 215.1, pp. 173–192.

Johri, Parul et al. (2022). “Recommendations for improving statistical inference in population genomics”. In: PLoS biology 20.5, e3001669.

Kant, Lakshmi, Shephalika Amrapali, and Banisetti Kalyana Babu (2016). “3 - Barley”. In: Genetic and Genomic Resources for Grain Cereals Improvement. Ed. by Mohar Singh and Hari D. Upadhyaya. San Diego: Academic Press, pp. 125–157. isbn: 978-0-12-802000-5. doi: 10.1016/B978-0-12-802000-5.00003-4. url: https://www.sciencedirect.com/science/article/pii/B9780128020005000034.

Kingman, J.F.C. (1982). “The coalescent”. In: Stochastic Processes and their Applications 13.3, pp. 235–248. issn: 0304-4149. doi: 10.1016/0304-4149(82)90011-4. url:https://www.sciencedirect.com/science/article/pii/0304414982900114.

Korfmann, Kevin et al. (2024a). “Determinants of rapid adaptation in species with large variance in offspring production”. In: Molecular Ecology 33.10, e16982.

Korfmann, Kevin et al. (2024b). “Simultaneous Inference of Past Demography and Selection from the Ancestral Recombination Graph under the Beta Coalescent”. en. In: Peer Community Journal 4, e33. doi: 10.24072/pcjournal.397. url: https://peercommunityjournal.org/articles/10.24072/pcjournal.397/.

Koskela, Jere (2018). “Multi-locus data distinguishes between population growth and multiple merger coalescents”. In: Statistical applications in genetics and molecular biology 17.3, p. 20170011.

Koskela, Jere and Maite Wilke Berenguer (2019). “Robust model selection between population growth and multiple merger coalescents”. In: Mathematical biosciences 311, pp. 1–12.

Koskela, Jere and Maite Wilke Berenguer (2019). “Robust model selection between population growth and multiple merger coalescents”. In: Mathematical Biosciences 311, pp. 1–12. issn: 0025-5564. doi: 10.1016/j.mbs.2019.03.004. url: https://www.sciencedirect.com/science/article/pii/S0025556418303523.

Kunz, Lukas, et al. (Feb. 2023). “The broad use of the Pm8 resistance gene in wheat resulted in hypermutation of the AvrPm8 gene in the powdery mildew pathogen”. In: BMC Biology 21.1. Publisher: Springer Science and Business Media LLC. issn: 1741-7007. doi: 10.1186/s12915-023-01513-5. url: https://bmcbiol.biomedcentral.com/articles/10.1186/s12915-023-01513-5.

Li, Heng (Nov. 2011). “A statistical framework for SNP calling, mutation discovery, association mapping and population genetical parameter estimation from sequencing data”. In: Bioinformatics 27.21, pp. 2987–2993.

Li, Heng (Sept. 2018). “Minimap2: pairwise alignment for nucleotide sequences”. In: Bioinformatics 34.18, pp. 3094–3100. issn: 1367-4803. doi: 10.1093/bioinformatics/bty191. eprint: https://academic.oup.com/bioinformatics/article-pdf/34/18/3094/48919122/bioinformatics_34_18_3094.pdf. url: https://doi.org/10.1093/bioinformatics/bty191.

Liu, Xinyi et al. (2026). Proteogenomics of Blumeria hordei supports RNA and protein coding innovative potential derived from transposable elements.

Matuszewski, Sebastian, et al. (Jan. 2018). “Coalescent Processes with Skewed Offspring Distributions and Nonequilibrium Demography”. In: Genetics 208.1, pp. 323–338. issn: 1943-2631. doi: 10.1534/genetics.117.300499. eprint: https://academic.oup.com/genetics/article-pdf/208/1/323/42168366/files4.pdf. url: https://doi.org/10.1534/genetics.117.300499.

McKenna, Aaron, et al. (Sept. 2010). “The Genome Analysis Toolkit: a MapReduce framework for analyzing next-generation DNA sequencing data”. en. In: Genome Res. 20.9, pp. 1297–1303.

Menardo, Fabrizio, Sébastien Gagneux, and Fabian Freund (Jan. 2021). “Multiple Merger Genealogies in Outbreaks of Mycobacterium tuberculosis”. In: Molecular Biology and Evolution 38.1, pp. 290–306. issn: 1537-1719. doi: 10.1093/molbev/msaa179. eprint: https://academic.oup.com/mbe/article-pdf/38/1/290/35389068/msaa179.pdf. url: https://doi.org/10.1093/molbev/msaa179.

Montano, Valeria (June 2016). “Coalescent inferences in conservation genetics: should the exception become the rule?” In: Biology Letters 12.6, p. 20160211. issn: 1744-9561. doi: 10.1098/rsbl.2016.0211. eprint: https://royalsocietypublishing.org/rsbl/article-pdf/doi/10.1098/rsbl.2016.0211/303012/rsbl.2016.0211.pdf. url: https://doi.org/10.1098/rsbl.2016.0211.

Müller, Marion C., et al. (Mar. 2019). “A chromosome-scale genome assembly reveals a highly dynamic effector repertoire of wheat powdery mildew”. In: New Phytologist 221.4. Publisher: Wiley, pp. 2176–2189. issn: 0028-646X, 1469-8137. doi: 10.1111/nph.15529. url: https://nph.onlinelibrary.wiley.com/doi/10.1111/nph.15529.

Papamakarios, George and Iain Murray (2016). “Fast *ɛ*-free Inference of Simulation Models with Bayesian Conditional Density Estimation”. In: Advances in Neural Information Processing Systems. Ed. by D. Lee et al. Vol. 29. Curran Associates, Inc. url: https://proceedings.neurips.cc/paper_files/paper/2016/file/6aca97005c68f1206823815f66102863-Paper.pdf.

Pavlidis, Pavlos, et al. (Sept. 2013). “SweeD: Likelihood-Based Detection of Selective Sweeps in Thousands of Genomes”. In: Molecular Biology and Evolution 30.9, pp. 2224–2234. issn: 0737-4038. doi: 10.1093/molbev/mst112. eprint: https://academic.oup.com/mbe/article-pdf/30/9/2224/13175153/mst112.pdf. url: https://doi.org/10.1093/molbev/mst112.

Pudlo, Pierre, et al. (Nov. 2015). “Reliable ABC model choice via random forests”. In: Bioinformatics 32.6, pp. 859–866. issn: 1367-4803. doi: 10.1093/bioinformatics/btv684. eprint: https://academic.oup.com/bioinformatics/article-pdf/32/6/859/49018353/bioinformatics_32_6_859.pdf. url: https://doi.org/10.1093/bioinformatics/btv684.

Sagitov, Serik (1999). “The General Coalescent with Asynchronous Mergers of Ancestral Lines”. In: Journal of Applied Probability 36.4, pp. 1116–1125. issn: 00219002. url: http://www.jstor.org/stable/3215582.

Schiffels, Stephan and Ke Wang (2020). “MSMC and MSMC2: The Multiple Sequentially Markovian Coalescent”. en. In: Methods Mol. Biol. Methods in molecular biology (Clifton, N.J.) 2090, pp. 147–166.

Schweinsberg, Jason (2003). “Coalescent processes obtained from supercritical Galton–Watson processes”. In: Stochastic Processes and their Applications 106.1, pp. 107–139. issn: 0304-4149. doi: 10.1016/S0304-4149(03)00028-0. url: https://www.sciencedirect.com/science/article/pii/S0304414903000280.

Sellinger, Thibaut Paul Patrick, Diala Abu-Awad, and Aurélien Tellier (2021). “Limits and convergence properties of the sequentially Markovian coalescent”. In: Molecular ecology resources 21.7, pp. 2231–2248.

Sotiropoulos, Alexandros G. et al. (July 2022). “Global genomic analyses of wheat powdery mildew reveal association of pathogen spread with historical human migration and trade”. In: Nature Communications 13.1. Publisher: Springer Science and Business Media LLC. issn: 2041-1723. doi: 10.1038/s41467-022-31975-0. url: https://www.nature.com/articles/s41467-022-31975-0.

Spanu, Pietro D. and Ralph Panstruga (2012). “Powdery mildew genomes in the crosshairs”. In: New Phytologist 195.1, pp. 20–22. doi: 10.1111/j.1469-8137.2012.04173.x. eprint: https://nph.onlinelibrary.wiley.com/doi/pdf/10.1111/j.1469-8137.2012.04173.x. url: https://nph.onlinelibrary.wiley.com/doi/abs/10.1111/j.1469-8137.2012.04173.x.

Spanu, Pietro D., et al. (Dec. 2010). “Genome Expansion and Gene Loss in Powdery Mildew Fungi Reveal Tradeoffs in Extreme Parasitism”. In: Science 330.6010. Publisher: American Association for the Advancement of Science (AAAS), pp. 1543–1546. issn: 0036-8075, 1095-9203. doi: 10.1126/science.1194573. url: https://www.science.org/doi/10.1126/science.1194573.

Stanca, A.M. et al. (2016). “Barley: An Overview of a Versatile Cereal Grain with Many Food and Feed Uses”. In: Reference Module in Food Science. Elsevier. isbn: 978-0-08-100596-5. doi: 10.1016/B978-0-08-100596-5.00021-4. url: https://www.sciencedirect.com/science/article/pii/B9780081005965000214.

Stephan, Wolfgang (2010). “Genetic hitchhiking versus background selection: the controversy and its implications”. In: Philosophical Transactions of the Royal Society B: Biological Sciences 365.1544, pp. 1245–1253.

Stephan, Wolfgang (2019). “Selective sweeps”. In: Genetics 211.1, pp. 5–13.

Talts, Sean et al. (2020). Validating Bayesian Inference Algorithms with Simulation-Based Calibration. arXiv: 1804.06788 [stat.ME]. url: https://arxiv.org/abs/1804.06788.

Tejero-Cantero, Alvaro et al. (2020). “sbi: A toolkit for simulation-based inference”. In: Journal of Open Source Software 5.52, p. 2505. doi: 10.21105/joss.02505. url: https://doi.org/10.21105/joss.02505.

Tellier, Aurélien and Christophe Lemaire (2014). “Coalescence 2.0: a multiple branching of recent theoretical developments and their applications”. In: Molecular Ecology 23.11, pp. 2637–2652. doi: 10.1111/mec.12755. eprint: https://onlinelibrary.wiley.com/doi/pdf/10.1111/mec.12755. url: https://onlinelibrary.wiley.com/doi/abs/10.1111/mec.12755.

VanLiere, Jenna M. and Noah A. Rosenberg (2008). “Mathematical properties of the r2 measure of linkage disequilibrium”. In: Theoretical Population Biology 74.1, pp. 130–137. issn: 0040-5809. doi: 10.1016/j.tpb.2008.05.006. url: https://www.sciencedirect.com/science/article/pii/S0040580908000609.

Vasimuddin, Md. et al. (2019). “Efficient Architecture-Aware Acceleration of BWA-MEM for Multicore Systems”. In: 2019 IEEE International Parallel and Distributed Processing Symposium (IPDPS), pp. 314–324. doi: 10.1109/IPDPS.2019.00041.

Wakeley, J. (Feb. 2009). “Coalescent Theory: An Introduction”. In: Systematic Biology 58. doi:10.1093/schbul/syp004.

Waples, Robin S (2016). “Tiny estimates of the N e/N ratio in marine fishes: Are they real?” In: Journal of Fish Biology 89.6, pp. 2479–2504.

Waples, Robin S et al. (2013). “Simple life-history traits explain key effective population size ratios across diverse taxa”. In: Proceedings of the Royal Society B: Biological Sciences 280.1768, p. 20131339.

Waples, Robin S et al. (2018). “Robust estimates of a high N e/N ratio in a top marine predator, southern bluefin tuna”. In: Science advances 4.7, eaar7759.

